# Evolutionary and functional genomics of DNA methylation in maize domestication and improvement

**DOI:** 10.1101/2020.03.13.991117

**Authors:** Gen Xu, Jing Lyu, Qing Li, Han Liu, Dafang Wang, Mei Zhang, Nathan M. Springer, Jeffrey Ross-Ibarra, Jinliang Yang

## Abstract

DNA methylation is a ubiquitous chromatin feature — in maize, more than 25% of cytosines in the genome are methylated. Recently, major progress has been made in describing the molecular mechanisms driving methylation, yet variation and evolution of the methylation landscape during maize domestication remain largely unknown. Here we leveraged whole-genome sequencing (WGS) and whole genome bisulfite sequencing (WGBS) on populations of modern maize, landrace, and teosinte (*Zea mays* ssp. *parviglumis*) to investigate the adaptive and phenotypic consequences of methylation variations in maize. By using a novel estimation approach, we inferred the methylome site frequency spectrum (mSFS) to estimate forward and backward epimutation rates and selection coefficients. We found weak evidence for direct selection on DNA methylation in any context, but thousands of differentially methylated regions (DMRs) were identified in population-wide that are correlated with recent selection. Further investigation revealed that DMRs are enriched in 5’ untranslated regions, and that maize hypomethylated DMRs likely helped rewire distal gene regulation. For two trait-associated DMRs, *vgt1*-DMR and *tb1*-DMR, our HiChIP data indicated that the interactive loops between DMRs and respective downstream genes were present in B73, a modern maize line, but absent in teosinte. Functional analyses suggested that these DMRs likely served as *cis*-acting elements that modulated gene regulation after domestication. Our results enable a better understanding of the evolutionary forces acting on patterns of DNA methylation and suggest a role of methylation variation in adaptive evolution.

## INTRODUCTION

Genomic DNA is tightly packed in the nucleus and is functionally modified by various chromatin marks such as DNA methylation of cytosine residues. DNA methylation is a heritable covalent modification prevalent in most species, from bacteria to humans [1, 2]. In mammals, DNA methylation commonly occurs in the symmetric CG context with exceptions of non-CG methylation in specific cell types, such as embryonic stem cells [3], but in plants it occurs in all contexts including CG, CHG and CHH (H stands for A, T, or C). Genome-wide levels of cytosine methylation exhibit substantial variation across angiosperms, largely due to differences in the genomic composition of transposable elements [4, 5], but broad patterns of methylation are often conserved within species [6, 7]. Across plant genomes, levels of DNA methylation vary widely from euchromatin to heterochromatin, driven by the different molecular mechanisms for the establishment and maintenance of DNA methylation in CG, CHG, and CHH contexts [8, 9].

DNA methylation is considered essential to suppress the activity of transposons [10], to regulate gene expression [11], and to maintain genome stability [8]. Failure to maintain patterns of DNA methylation in many cases can lead to developmental abnormalities and even lethality [12–14]. Nonetheless, variation in DNA methylation has been detected both in natural plant [15] and human populations [16]. Levels of DNA methylation can be affected by genetic variation and environmental cues [17]. Additionally, heritable *de novo* epimutation — the stochastic loss or gain of DNA methylation — can occur spontaneously and has functional consequences [18, 19]. Population methylome studies suggest that the spread of DNA methylation from transposons into flanking regions is one of the major sources of epimutation, such that 20% and 50% of the *cis*-meQTL (methylation Quantitative Trait Loci) are attributable to flanking structural variants in *Arabidopsis* [7] and maize [20].

In *Arabidopsis,* a multi-generational epimutation accumulation experiment [21] estimated forward (gain of DNA methylation) and backward (loss of methylation) epimutation rates per CG site at about 2.56 × 10^−4^ and 6.30 × 10^−4^, respectively. Other than this *Arabidopsis* experiment, there are no systematic estimates of the epimutation rates in higher plants (but see recently estimates for poplar and dandelion [22]), making it difficult to understand the extent to which spontaneous epimutations contribute to methylome diversity in a natural population. Because the per base rates of DNA methylation variation are several orders of magnitude larger than DNA point mutation, conventional population genetic models which assume infinite sites models seemed inappropriate for epimutation modeling. As an attempt to overcome the obstacle, Charlesworth and Jain [23] developed an analytical framework to address evolution questions for epimutations. Leveraging this theoretical framework, Vidalis et al. [24] constructed the methylome site frequency spectrum (mSFS) using worldwide *Arabidopsis* samples, but they failed to find evidence for selection on genic CG epimutation under benign environments. The confounding effect between DNA variation and methylation variation, as well as the high scaled epimutation rates become obstacles to further dissect the evolutionary forces in shaping the methylation patterns at different timescales under different environments.

Maize, a major cereal crop species, was domesticated from its wild ancestor teosinte (Z. *mays* ssp. *parviglumis*) near the Balsas River Valley area in Mexico about 9,000 years ago. Genetic studies reveal that the dramatic morphological differences between maize and teosinte are largely due to selection of several major effect loci [25]. As maize spread across the Americas, many additional loci have played an important role in local adaptation [26]. Flowering time, a trait that directly affects plant fitness, played a major role in this local adaptation process [27–29]. Previous research, however, has focused almost entirely on DNA variation, and the contributions of methylation variation to maize domestication and adaptation remain largely elusive.

Here, we collected a set of geographically widespread Mexican landraces and a natural population of teosinte near Palmar Chico, Mexico [30], from which we generated genome and methylome sequencing data. Additionally, we profiled the teosinte interactome using HiChIP. Together with the analysis from previously published genome [31], transcriptome [32], methylome [6], and interactome [33] datasets, we estimated epimutation rates and selection pressures across different timescales, investigated the DNA methylation landscape in maize and teosinte, detected differentially methylated regions (DMRs), characterized the genomic features that are related with DMRs, and functionally validated two DMRs that are associated with adaptive traits. Our results suggest that DNA methylation genome-wide is likely only under relatively weak selection, but that methylation differences at a subset of key loci may modulate the regulation of domestication genes and affect maize adaptation.

## RESULTS

### Genomic distribution of methylation in maize and teosinte

To investigate genome-wide methylation patterns in maize and teosinte, we performed whole-genome bisulfite sequencing from a panel of wild teosinte, domesticated maize landraces, and modern maize inbreds (**Table S1)**. Using the resequenced genome of each line, we created individual pseudo-references (see **methods**) that alleviated potential bias of mapping reads to a single reference genome [34] and improved overall read-mapping (**Figure S1A**). Using pseudo-references, on average about 25 million (5.6%) more methylated cytosine sites were identified than using the B73 reference (**Figure S1B**). Across populations, average genome-wide cytosine methylation levels were about 78.6%, 66.1% and 2.1% in CG, CHG, and CHH contexts, respectively, which are consistent with previous estimations in maize [13] and are much higher than observed (30.4% CG, 9.9% CHG, and 3.9% CHH) in *Arabidopsis* [5]. We observed slightly higher levels of methylation in landraces, which may be due to lower sequencing depth [35]. We found no significant differences between teosinte and maize as a group (**Figure S2)**.

We found methylated cytosines in CG and CHG contexts were significantly higher in pericentromeric regions (0.54 ± 0.01 in a 10 Mb window) than in chromosome arms (0.44 ± 0.04) (Students’ t test P-value < 2.2e – 16) (**Figure S3)**. We calculated the average methylated CG (mCG) level across gene bodies (from transcription start site to transcription termination site, including exons and introns) in each population and observed a bimodal distribution of mCG in gene bodies (**Figure S4A**), with approximately 25% of genes (N = 6, 874) showing evidence of gene body methylation (gbm). While the overall distribution of gbm did not differ across populations, genes with clear syntenic orthologs in *Sorghum* [36] exhibited little gbm (**Figure S4B-C**), consistent with previous reports [5, 37].

### Genome-wide methylation is only under weak selection

As the frequency of methylation may be affected by both selection and epimutation rates, we implemented a novel MCMC approach to estimate these parameters using a population genetic model developed for highly variable loci [23]. We defined 100-bp tiles across the genome as a DNA methylation locus and categorized individual tiles as unmethylated, methylated, or heterozygous alleles for outcrossed populations (i.e., teosinte and landrace populations) and as unmethylated and methylated alleles for modern maize inbred lines (see **methods**). To determine the thresholds for methylation calls, we employed an iterative expectation maximization algorithm to fit the data [38]. We then constructed methylome site frequency spectra (mSFS) for CG and CHG sites (**Figure S5)**. Sensitivity test results suggested that the mSFS was insensitive to the cutoffs used for the methylation calls (**Figure S6)**. As the vast majority (≥ 98%) of CHH sites were unmethylated (**Figure S7)**, we excluded CHH sites from population genetic analysis.

After testing a set of prior values, we found the initial prior rates had little impact on the posteriors, except for extremely large values (**Figure S8)**, for which convergence was difficult. Because we found little difference among populations in genome-wide patterns, we estimated parameters using the combined data; estimates from individual populations were nonetheless broadly similar (**Figure S9)**. Effective population size (*N_e_*) in maize is difficult to estimate because of rapid demographic change during and post-domestication. Previous estimates of *N_e_* in maize range from ~ 50k [39] to ~ 370*k* – 1*M* [40]. To account for this uncertainty, we ran the models with a set of different *N_e_* values (50*k*, 100*k*, 500*k*, and 1*M*). Model estimates of the epimutation rate *μ* for both CG (3.6 × 10^−6^ – 1.8 × 10^−7^) and CHG (7.6 × 10^−6^ – 3.8 × 10^−7^) sites were more than an order of magnitude higher than the backward epimutation rates (*v* = 1.8 × 10^−7^ – 9.0 × 10^−9^ and 3.0 × 10^−7^ – 1.5 × 10^−8^) using different *N_e_* values (**Figure 1A**), consistent with the observed prevalence of both types of methylation. Estimates of the genome-wide selection coefficient *s* associated with methylation of a 100-bp tile for both CG and CHG tiles depended on the assumption of *N_e_.* However, the population-scaled selection coefficient (or *N_e_* × *s*) stayed largely constant with values of 2.0 and 2.2 for CG and CHG tiles, indicating relatively weak selection for methylation in each context according to classical population genetic theory [41].

**Fig. 1.**
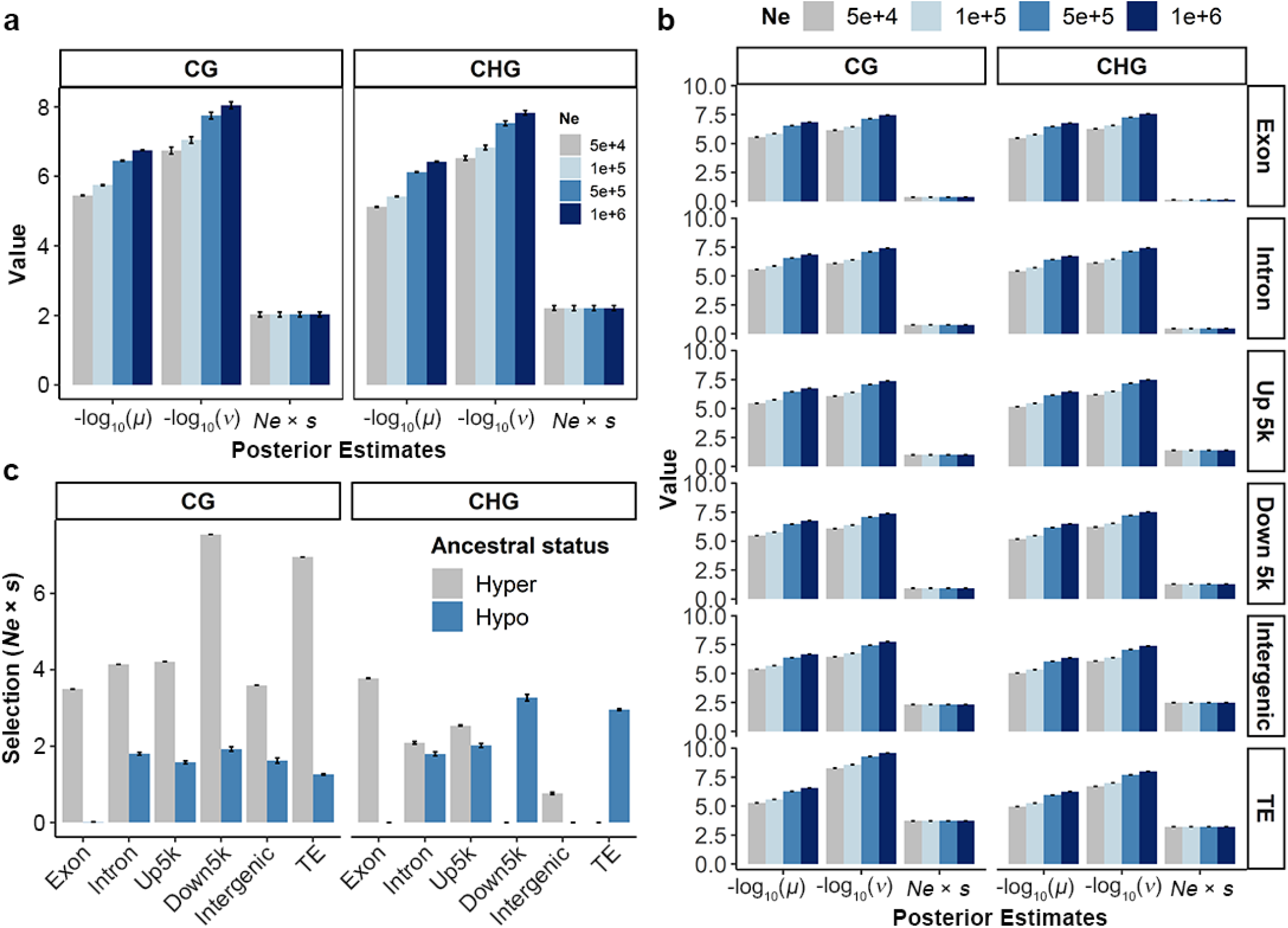
Population genetic parameters inference. (**a**) Posterior estimates of mean values and standard deviations for *μ, v,* and *N_e_* × *s* for CG and CHG sites using four different effective population size (*N_e_*) values. (**b**) Posterior estimates for different genomic features. Up 5k, the upstream 5k region of a gene; Down 5k, the downstream 5k region of a gene. (**c**) Posterior estimates by defining teosinte as the ancestral epiallele. Values were estimated using MCMC approach with 20% burnin (see **methods**). Error bars indicate standard deviations. Source Data underlying (**a**-**c**) are provided in a Source Data file. (N = 1,600 for each bar).

We then sought to test whether the population-scaled selection coefficient differs across genomic features. After fitting mSFS models separately for different genomic features, results showed that population-scaled selection coefficients in genic regions (exon, intron, upstream 5k, and downstream 5k) were below or close to 1, and the values were above 1 for non-genic regions (i.e., 2.4 for intergenic regions and 3.5 for TE regions) (**Figure 1B**), suggesting stronger selection on methylation variation outside of genes. If we consider the most common variant in teosinte as the ancestral epiallele, selection was higher in ancestrally hypermethylated regions in CG contexts, especially in TE and intergenic regions, while it was close to neutrality for ancestrally hypomethylated regions, especially for the exonic regions (**Figure 1C**). In CHG contexts, selection was weak in most regions, including TE and intergenic regions, for both ancestral hyper- and hypomethylated sites.

### Regions with variable methylation contribute to phenotypic variation

Our observed CG mSFS revealed that 2% and 7% of 100-bp tiles were completely unmethylated and methylated, while 91% of tiles were variable (**Figure S5A**). These variable methylation regions can be further divided into rarely unmethylated (frequency of methylated tiles > 90%), rarely methylated (frequency of methylated tiles < 10%), and high-frequency variable regions (frequency of methylated tiles >= 10% and <= 90%), composing 69%, 2%, and 20% of the genome, respectively. To investigate whether regions of the genome exhibiting variable methylation, especially the high-frequency variable regions, are functionally relevant, we used SNP sets residing in these five regions to estimate their contribution to phenotypic variation using published data from a large maize mapping population [42]. We estimated kinship matrices for SNPs in different genomic regions and then partitioned the genetic variance for plant phenotypes using LDAK [43]. Consistent with an important functional role for genic regions and a lack of functional importance in permanently methylated regions, our results find that sites that are hypomethylated (uniformly unmethylated and rarely methylated), mainly from the genic areas, explained disproportionally larger genetic variances (**Figure S10A**). While hypermethylated regions (uniformly methylated and rarely unmethylated), although accounting for 76% of the genome, contribute only a fraction of the genetic variance for 7/23 traits. The proportion of variance explained by high-frequency sites polymorphic for methylation, ranged from 0% to 57%, with a mean value of 29%. Variance component analysis results for CHG sites were largely consistent with the results for CG sites (see **Figure S10B**).

### Population level DMRs are enriched in selective sweeps

Although genome-wide selection on epimutation appears relatively weak, the observation that sites exhibiting methylation polymorphism contribute meaningfully to quantitative trait variation suggested that stronger selection could be acting at specific differentially methylated regions (DMRs). We employed the metilene software [44] to identify a total of 5,278 DMRs (see **Table 1** for numbers broken down by context and type), or about 0.08% (1.8 Mb) of the genome, including 3,900 DMRs between teosinte and modern maize, 1,019 between teosinte and landrace, and 359 DMRs between landrace and modern maize (**Table S2)**. To check the tissue-specificity of the detected DMRs, we examined the methylation levels of these DMRs in B73 across different tissue types using published WGBS data [45]. Our results suggested that methylation levels of the DMRs were largely conserved in B73 across three tissue types (**Figure S11)**, consistent with the previous studies [20, 46, 47].

**Table 1.**
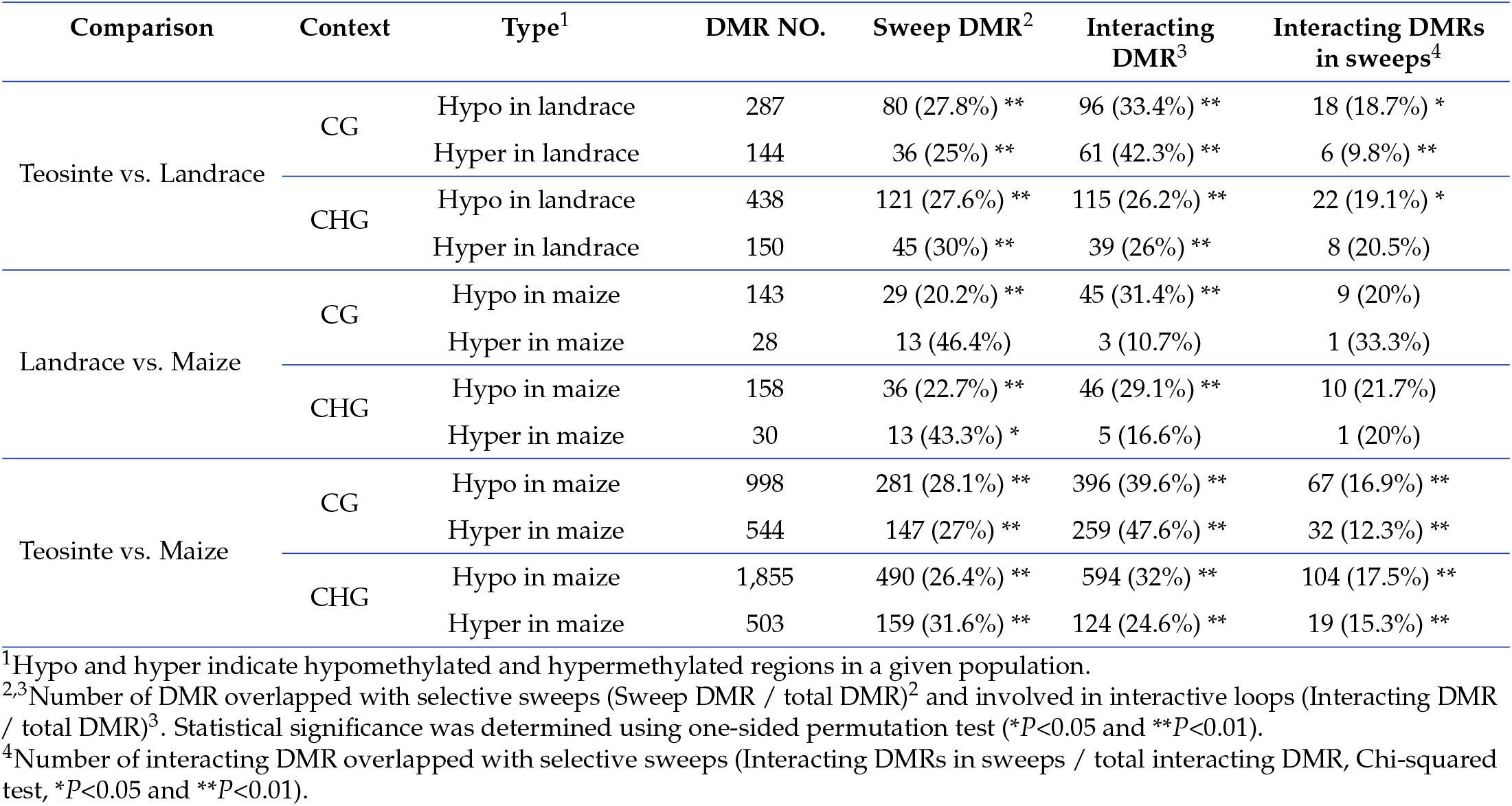
Number of differentially methylated regions (DMRs) broken down by context and type.

DNA methylation can have a number of functional consequences [15, 48, 49], and thus we tested whether differences in methylation among populations were associated with selection at individual locus. To test this hypothesis, we used SNP data from each population to scan for genomic regions showing evidence of selection (see **methods**). We detected a total of 1,330 selective sweeps between modern maize and teosinte (**Figure 2** and **Table S3,** see **Figure S12** for results of teosinte vs. landrace and landrace vs. modern maize). Several classical domestication genes, e.g., *tb1* [50], *ZAG2* [51], *ZmSWEET4c* [52], *RA1* [53], and *BT2* [54] were among these selective signals.

**Fig. 2.**
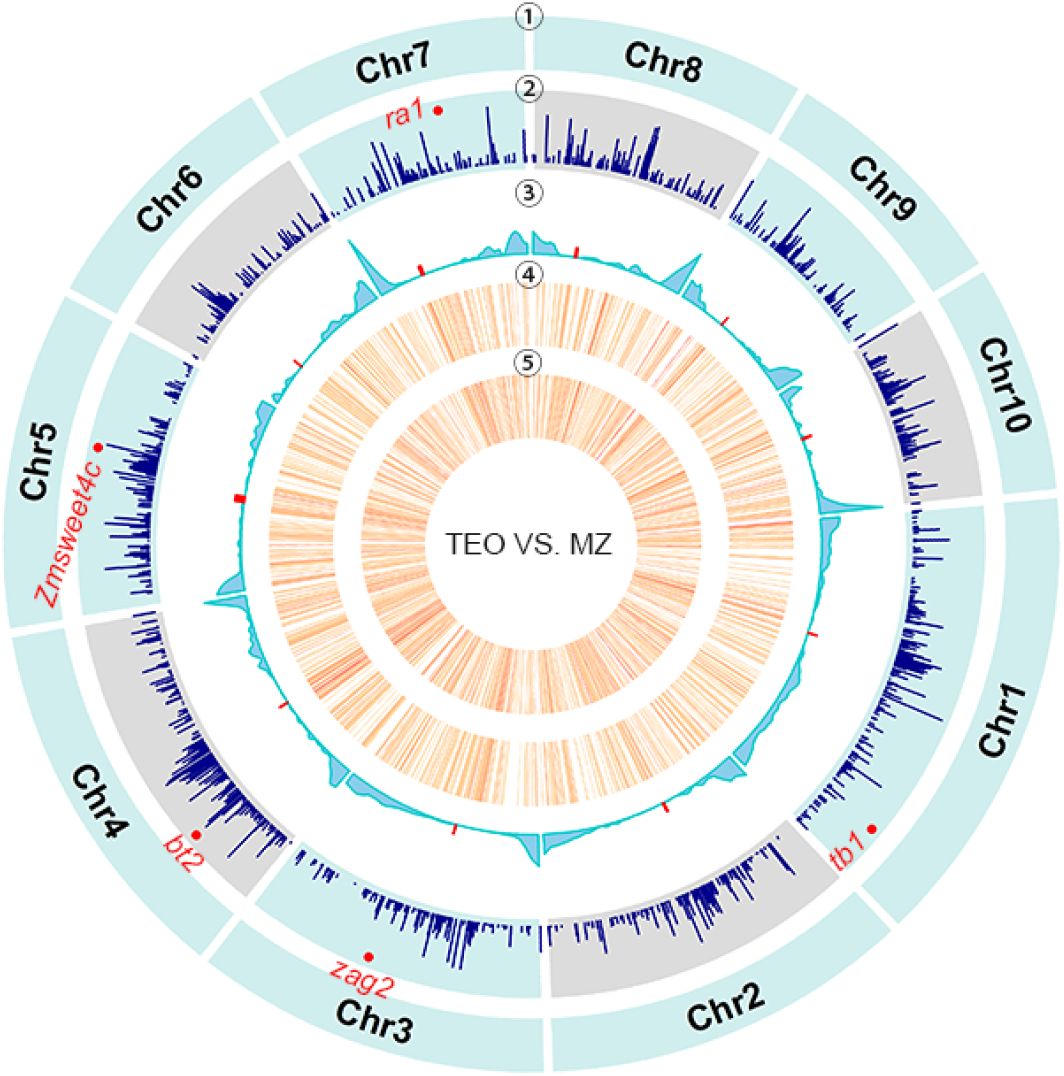
Selection on differentially methylated regions. Distributions of teosinte-maize selective sweeps, DMRs and other genomic features across ten maize chromosomes. From outer to inner circles were ① Chromosome names, ② selective sweeps detected between modern maize and teosinte, ③ the recombination rate, and the density of DMRs (number per 1-Mb) between modern maize and teosinte in ④ CG and ⑤ CHG contexts. Red dots in circle ③ denote the centromeres. Source Data underlying are provided in a Source Data file.

We found that DMRs at CG and CHG sites were highly enriched in regions showing evidence of recent selection (**Figure S13,** P-value < 0.001), particularly in intergenic and TE regions (**Figure S14A**). DMRs overlapping sweeps, both hypo- and hypermethylated in maize, exhibited significantly higher allele frequency differentiation between maize and teosinte (**Figure S14B**, see **Table 1** for other comparisons). We then asked whether DMRs in sweep regions were in linkage disequilibrium with nearby SNPs (see **methods**), as might be expected if most DMRs were the result of an underlying genetic change such as a TE insertion. Indeed, the rate of sweep DMRs in LD with local SNPs was significantly higher than expected by chance (**Table S4)**.

Additionally, we detected 72 genes located in sweep DMRs (maize vs. teosinte under CG context) that were hypomethylated in maize, 24 (42/72 with expression data) of which showed significantly (Student’s paired t-test, P-value < 0.05) increased expression levels in maize compared to teosinte using published data [32]. For the 56 genes located in sweep DMRs that were hypermethylated in maize, however, we failed to detect the significant expression differences between maize and teosinte.

### Hypomethylated regions in maize are associated with interacting loops

Further investigation indicated that teosinte-maize CG DMRs were significantly enriched in mappable genic and intergenic (i.e., nongenic excluding 5-kb upstream and downstream of genes and transposons) regions for both hyper- and hypomethylated regions in maize, but depleted from transposon regions (**Figure 3A**). We detected maize hyper- and hypomethylated DMRs in 0.01% and 0.02% of mappable regions across the genome. In particular, 0.07% and 0.05% of maize hyper-DMR (DMR hypermethylated in maize) and hypo-DMR (DMR hypomethylated in maize) were located within mappable exonic regions, which were much higher than expected by chance (permutation P-values < 0.001, **Figure S15A**). These CG DMRs could be mapped to *N =* 229 unique genes (**Table S5)**. After examining the mapping locations based on collapsed gene models, we found that DMRs were most abundant in at 5’ UTRs (**Figure 3B**), consistent with a pattern that was previously observed [56]. Using these DMR genes for a gene ontology (GO) analysis, we detected 15 molecular function terms that were significantly enriched (**Figure S15B**). The vast majority (14/15) of these significant terms were associated with “binding” activities, including protein, nucleoside, and ribonucleoside binding. Furthermore, we found that exonic DMRs were enriched at transcription factor binding sites identified via DAP-seq [57] (permutation P-value < 0.001).

**Fig. 3.**
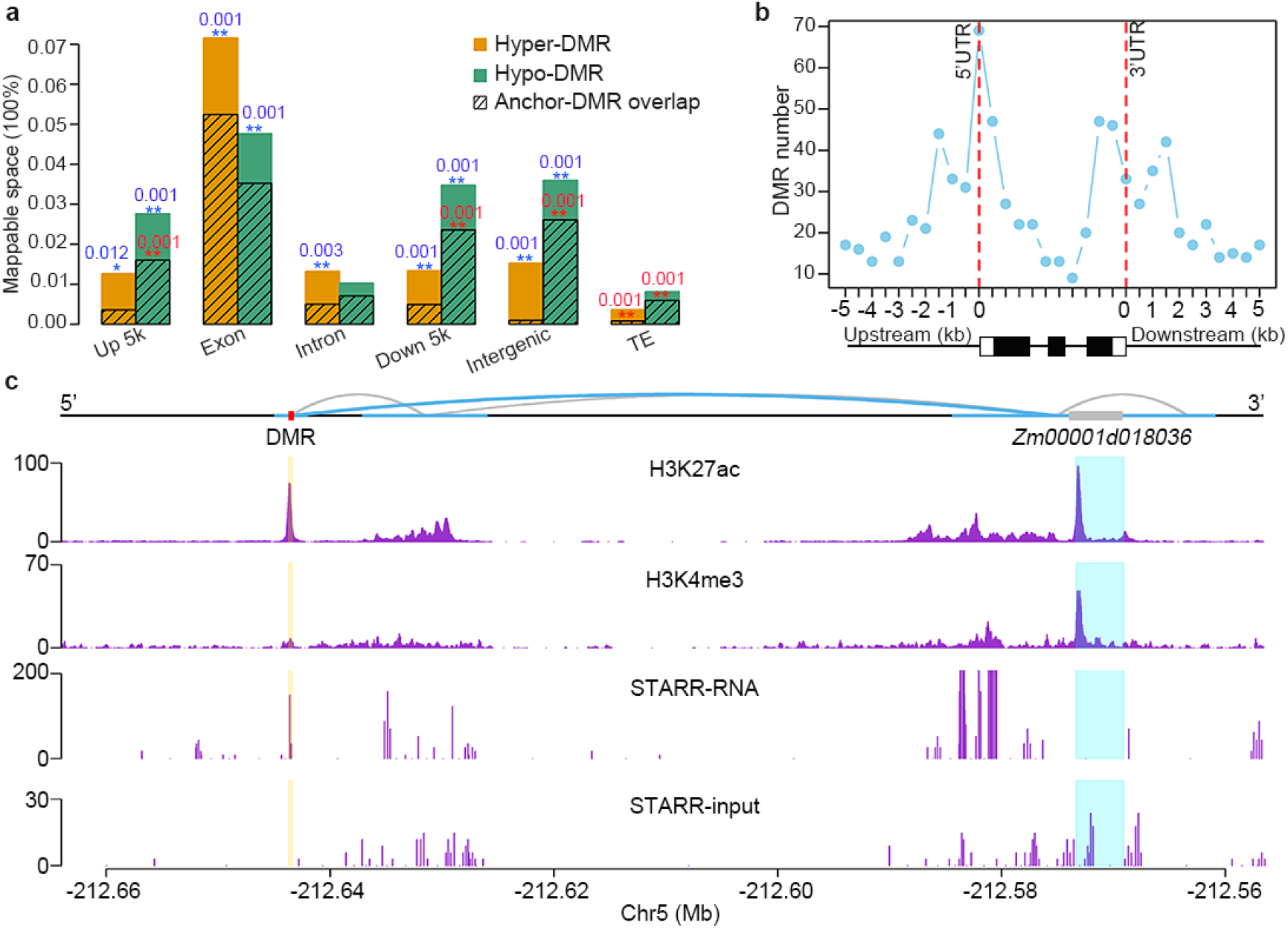
Teosinte-maize CG DMRs and their associated functional features. (**a**) Breakdown of hyper-DMRs (DMR hypermethylated in maize) and hypo-DMRs (DMR hypomethylated in maize) into genomic features and their overlaps with interactive anchors using data obtained from Li et al., [33]. Blue and red stars indicated DMRs that were significantly enriched at genomic features and interaction anchors (one-sided permutation test: *P-value < 0.05, *\*\*P*-value < 0.01). The numbers above the asterisks indicate the exact test P-values. (**b**) The distribution of the number of DMRs along the collapsed gene model. Below the figure shows a schematic gene model with three exons (black boxes). (**c**) Physical interactions (upper panel), colocalization with H3K27ac and H3K4me3 (middle panels), and STARR profiles (lower panels) around *Zm00001d018036* gene in B73. STARR-seq data obtained from [55] showed the transcriptional output (STARR-RNA) and DNA input (STARR-input) around this region. Blue curly lines indicate the interactive contacts between DMR and the candidate gene and grey curly lines indicate other interactive contacts around the region. Horizontal thick blue lines denote the interactive anchors. Red and grey boxes indicate the DMR and gene model, respectively. Source Data underlying (**a**, **b**) are provided in a Source Data file.

These findings suggested a potential role for DMRs affecting regulatory regions. To investigate this possibility, we made use of recent data using long-read ChIA-PET to profile genomic regions colocalized with H3K4me3 and H3K27ac to define the interactome in maize [33]. We found that interactive anchor sequences were significantly enriched in DMRs that are hypomethylated in maize, especially in regulatory regions, including upstream 5-kb, downstream 5-kb, and intergenic regions (**Figure 3A**). We also found that DMRs located in transposable elements that were hypomethylated in maize more likely overlap with interactive anchors than expected by chance (permutation *P*-value < 0.001).

We hypothesized that these hypomethylated DMRs, especially intergenic DMRs overlapped with the regulatory regions, will alter the up- or downstream gene expression through physical interactions. To test this hypothesis, we mapped the interactive anchors harboring maize hypomethylated DMRs to their 1st, 2nd, and 3rd levels of contacts (**Figure S16A**). Interestingly, among the 60 genes in direct contact with maize hypomethylated intergenic DMRs (**Table S6)**, we found that 30 (43/60 with expression data) showed significantly (Student’s paired t-test, P-value < 0.05) increased expression levels in maize compared to teosinte using published data [32]. The results were not significant for 2nd and 3rd level contacts (**Figure S16A**). We found 5/60 genes (Enrichment test P-value < 0.01) were domestication candidate genes as reported previously [58–61]. Two of them were *Zm00001d018036* (a gene associated with cob length, P-value = 6 × 10^-25^) and *Zm00001d041948* (a gene associated with shank length, P-value = 5.6 × 10^-10^) [58]. Further investigation of these two candidates using recently published chromatin data [55] to detect enhancer activity [62] identified H3K27ac peaks at both DMR loci (**Figure 3C** and **Figure S17A**). Consistent with these enhancer signals, the expression levels of these two genes is significantly increased in maize relative to teosinte (**Figure S16B** and **Figure S17B**). Despite this functional evidence, however, we found that interacting DMRs in selective sweeps were significantly less often than expected by chance (**Table 1)**.

### DMRs associated with flowering time variation

Analyses above found that high-frequency regions polymorphic for methylation in our samples accounted for 15% and 17% genetic variances for two flowering time traits, days to anthesis and days to silk, respectively (**Figure S10A**). Upon closer inspection of our DMRs, we found a number of candidate flowering time genes located in sweep DMRs or interacting DMRs (**Table S7)**, including three genes found in both (i.e., *Zm00001d029946, Zm00001d015884, Zm00001d025979).* We also examined several known genes in the flowering time pathway [63] and detected six DMRs located near four additional flowering time related genes (**Figure S18)** (Enrichment test P-value < 0.05).

One DMR was located 40-kb upstream of *ZmRAP2.7,* a well characterized flowering time gene, and 20-kb downstream of the *vgt1* locus, that was hypomethylated in modern maize and landrace but was hypermethylated in teosinte (**Figure 4A**). A MITE transposon insertion in the *vgt1* locus is considered as the causal variant for the down regulation of *ZmRAP2.7,* which encodes a transcription factor in the flowering time pathway [64]. We did not detect *vgt1* as a selective sweep because it is not considered a domestication or improvement candidate and our maize lines include both tropical and temperate lines [65]. We further examined LD in this regions and detected strong signals between the vgt1-DMR and local SNPs, suggesting that the vgt1-DMR is not a pure epiallele. Reanalysis of published ChIP data [33] revealed that the DMR colocalized with a H3K27ac peak and there is a physical interaction between the DMR and the *vgt1* locus in maize [33] (**Figure 4B**). Additionally, we reanalyzed the maize and sorghum sequence data at the *vgt1* locus and found two conserved non-coding sequences (CNSs) located 1kb downstream of the vgt1-DMR (**Figure S19)**. To examine the interaction status in teosinte, we then generated HiChIP data for a teosinte sample using the same tissue and antibodies (see **methods**). Although our teosinte HiChIP data identified similar peaks of H3K27ac and H3K4me3 near the region, we failed to detect a physical interaction between the *vgt1*-DMR and *vgt1* itself in teosinte (**Figure 4B**), suggesting that methylation at this locus might play a functional role in affecting physical interaction.

**Fig. 4.**
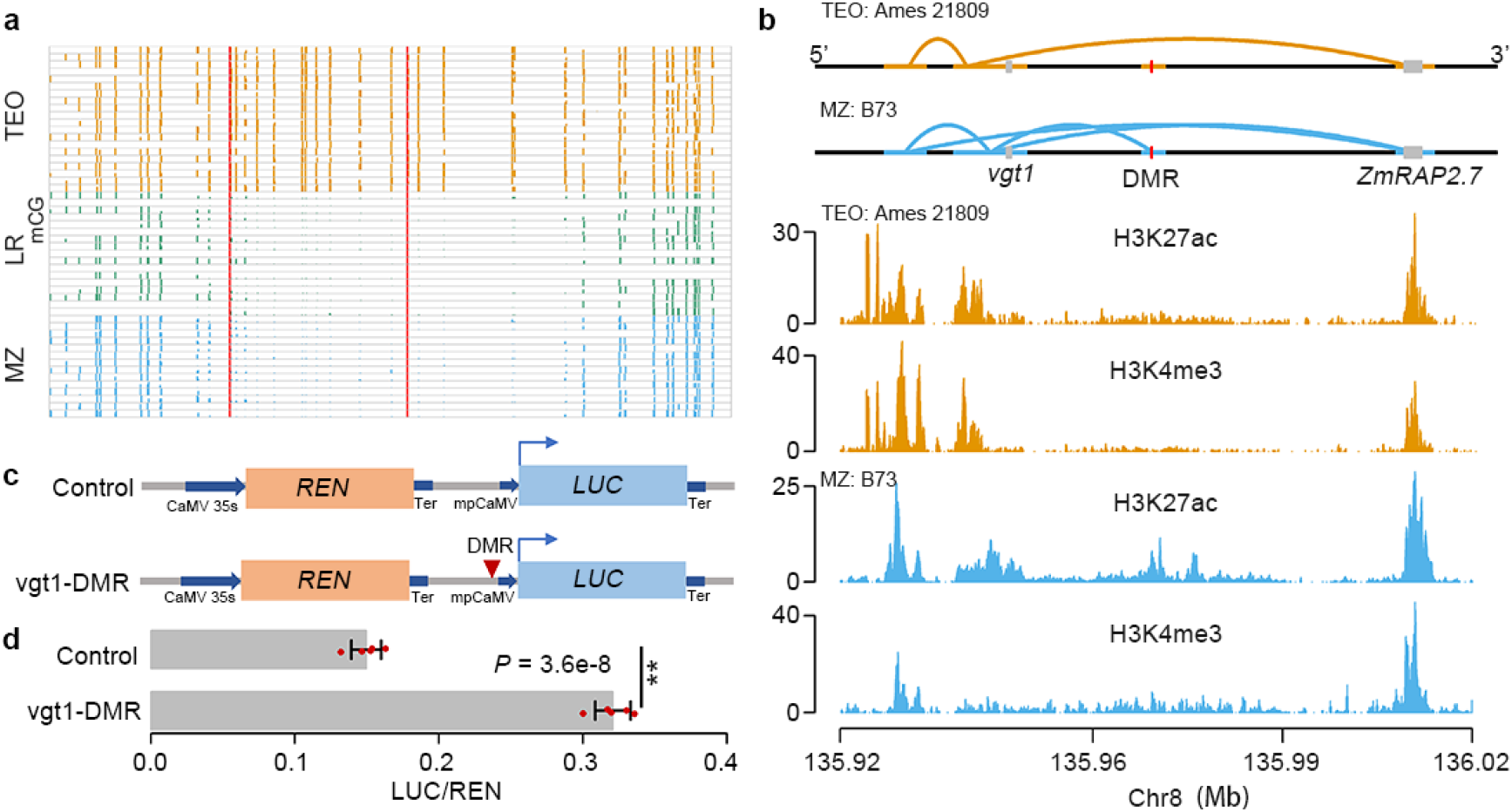
Functional analysis of vgt1-DMR. (**a**) Levels of CG methylation around *vgt1*-DMR in maize (MZ), landrace (LR), and teosinte (TEO) populations. Vertical red lines indicate the boundaries of the vgt1-DMR. (**b**) The interactive contacts (upper panel) and colocalization with H3K27ac and H3K4me3 (lower panel) around vgt1-DMR in a maize (B73) and a teosinte (Ames 21809) samples. (**c**) The vectors constructed for functional validation of the *vgt1*-DMR using the dual-luciferase transient expression assay in maize protoplasts. (**d**) The expression ratios of LUC/REN using five biological replicates. Error bars indicated standard deviations. Statistical significance was determined by a two-sided t-test (**P-value = 3.6e – 8). Source Data underlying (**a**, **d**) are provided in a Source Data file.

To further validate the potential enhancer function of the 209-bp *vgt1*-DMR, we incorporated the *vgt1*-DMR sequence amplified from B73 into a vector constructed as shown in (**Figure 4C**) and performed a dual-luciferase transient expression assay in maize protoplasts (see **methods**). The results of the transient expression assay revealed that the maize cells harboring the DMR exhibited a significantly higher LUC and REN ratio than control (fold change= 2.2, P-value= 2.4e^−8^, **Figure 4D**), revealing that the DMR might act as an enhancer to activate LUC expression.

### A segregating tb1-DMR acts like a *cis*-acting element

One of the most significant teosinte-maize CG DMRs was located 40-kb upstream of the *tb1* gene, which encodes a transcription factor acting as a repressor of axillary branching (aka tillering) phenotype [50]. This 534-bp *tb1*-DMR was hypomethylated in modern maize, hypermethylated in teosinte, and segregating in landraces (**Figure 5A**). Chop-PCR (DNA methylation-sensitive restriction endonuclease digestion followed by PCR) analysis using a modern maize (inbred line W22) and a teosinte accession (PI 8759) suggested that DNA methylation presents in both leaf and immature ear tissues in teosinte, but is absent in W22 (**Figure S20)**. The physical location of the tb1-DMR overlapped with the MNase hypersensitive site [66] and a H3K9ac peak [67]. Phenotypic analysis of our 17 landraces indicated that the DMR was associated with the tillering (Fisher’s exact test P-value < 0.05), consistent with previous observations that the hypermethylated (teosinte-like) genotypes were likely to grow tillers [50].

**Fig. 5.**
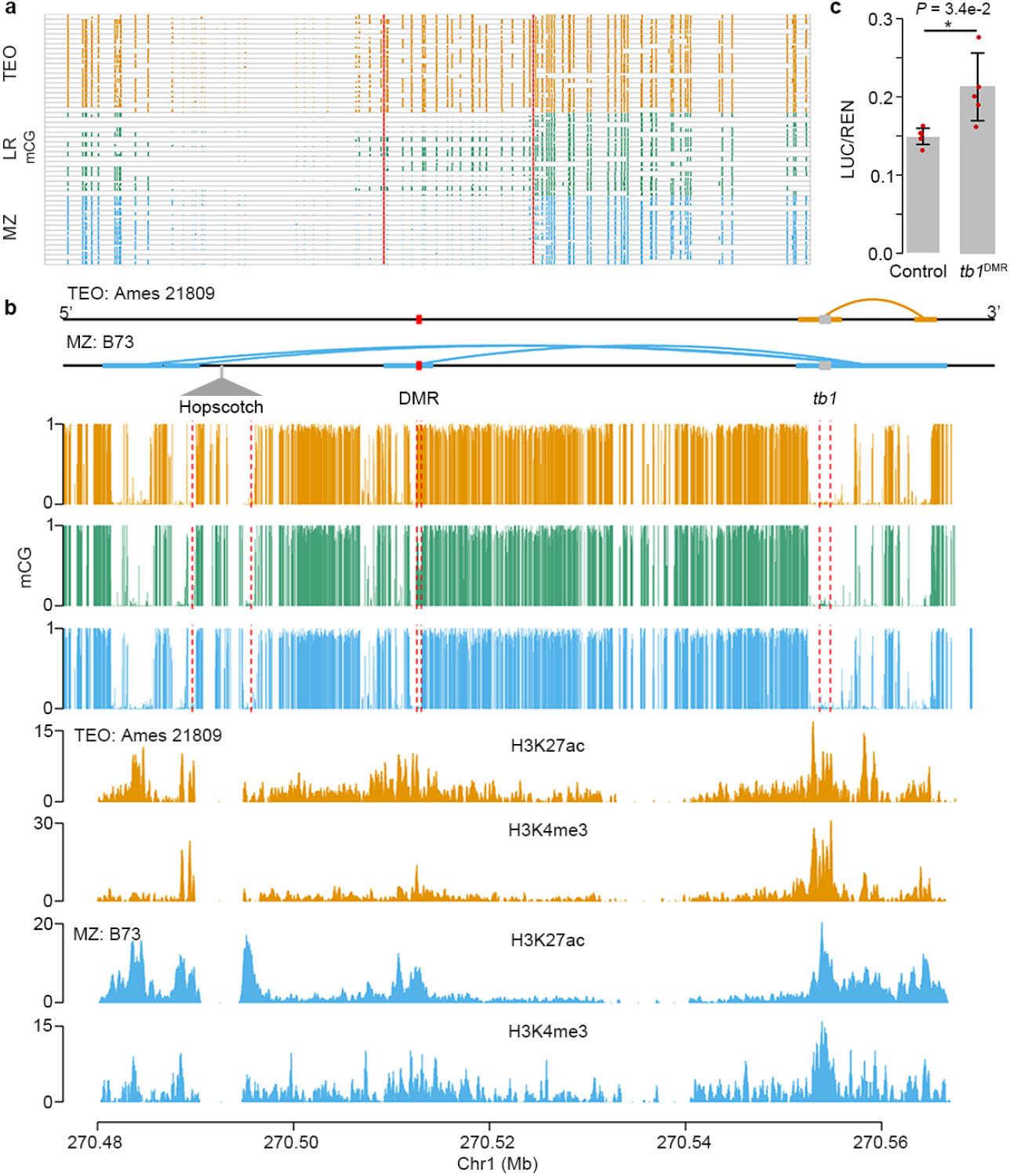
A hypomethylated DMR that is upstream of tb1 gene. (**a**) Levels of mCG for the 534-bp tb1-DMR in each individual methylome of the modern maize (MZ), landrace (LR), and teosinte (TEO) populations. Vertical red lines indicate the boundaries of the tb1-DMR. (**b**) Interactive contacts (upper panel), average CG methylation levels (middle panel), and colocalization of the tb1-DMR with H3k27ac and H3K4me3 (lower panel). Horizontal thick lines denote the interactive anchors and solid curly lines on top of the annotations denote the interactive contacts in teosinte and maize. (**c**) Functional validation result of tb1-DMR using dual-luciferase transient expression assay in maize protoplasts. Five biological replicates were performed. Error bars indicated standard deviations. Statistical significance was determined by a two-sided t-test (*P-value = 3.4e – 2). Source Data underlying (**a**, **c**) are provided in a Source Data file.

The causal variation for this locus was previously mapped to a Hopscotch TE insertion 60-kb upstream (**Figure 5B**) of the *tb1* gene. The TE was considered as an enhancer, as shown by a transient *in vivo* assay [50]. Interactome data support this claim, finding physical contact between Hopscotch and the *tb1* gene (**Figure 5B**) [33]. Direct physical contact between the *tb1*-DMR and the *tb1* gene itself in maize line B73 was also detected using ChIA-PET data [33], but this interaction is missing in teosinte based on our newly generated HiChIP data (**Figure 5B**). By employing the 4C-seq method [68], we further confirmed the absence of interaction between the *tb1*-DMR and the *tb1* gene using landrace samples showing hypermethylation at the *tb1*-DMR locus (**Figure S21)**. The colocalization of *tb1*-DMR with chromatin activation marks in the region also suggested the *tb1*-DMR might act as a *cis*-acting regulatory element (**Figure 5B**). Additionally, we conducted a dual-luciferase transient assay by constructing a vector similar to the vgt1-DMR (**Figure 4E**). The results indicated that the tb1-DMR significantly increased the LUC/REN ratio as compared to control (**Figure 5C**), suggesting that the *tb1*-DMR was potentially act as a *cis*-acting element to enhance downstream gene expression.

To understand the correlation among these genomic components, i.e., the tb1-DMR, the TE insertion, and the *tb1* gene, we conducted linkage disequilibrium (LD) analysis using landrace genomic and methylation data segregating at this *tb1*-DMR locus (see **methods**). As a result, we failed to detect strong LD (R^2^ = 0.1) between the tb1-DMR and SNPs located at the Hopscotch locus (**Figure S22)**, indicating the *tb1*-DMR might be independent from the Hopscotch locus. Reanalysis of published *tb1* mapping data [50] confirmed a significant QTL signal around the Hopscotch TE (**Figure S23A**), and a two-dimensional QTL scan detected epistasis between Hopscotch and the *tb1*-DMR (**Figure S23B**). Further, we found that highly methylated landraces were geographically closer to the Balsas River Valley in Mexico, where maize was originally domesticated from (**Figure S24A**). As the landraces spread out from the domestication center, their CG methylation levels were gradually reduced (**Figure S24B**).

## DISCUSSION

In this study, we employed population genetics and statistical genomics approaches to infer the rates of epimutation and selection pressure on DNA methylation, and the extent to which SNPs located within DMRs contributed to phenotypic variation. Our results revealed that the forward epimutation rate was about 10 times larger than the backward epimutation rate. These estimates from 100-bp tiles are lower than epimutation rates estimated at nucleotides in *Arabidopsis* from epimutation accumulation experiments [69]. Even so, our estimated epimutation rates are more than an order of magnitude higher than the per-nucleotide mutation rate in maize [70].

Although population methylome modeling suggested that genome-wide DNA methylation was not under strong selection, we nonetheless show that regions harboring polymorphic methylation contribute to functionally relevant phenotypic variation. To prioritize loci likely exhibiting evolutionarily relevant methylation variation, we identified individual differentially methylated regions (DMRs). These DMR were enriched in likely functional sequence, including regulatory regions near genes, putative enhancers, and intergenic regions showing evidence of chromatin interactions. We further identified several dozen new genes that are differentially expressed between maize and teosinte, for which exonic regions directly interact with maize hypo-DMRs. We also found a strong enrichment of DMRs in regions targeted by recent positive selection. Patterns of linkage disequilibrium between DMRs and nearby SNPs make it difficult to assign causality, i.e., the DMRs associated with the flowering time traits maybe not the causal variants, but are consistent with the idea that many DMRs are the result of genetic changes, consistent with previous studies [7, 20]. Taken together, these results suggest that methylation might modulate physical interactions and hence likely affect gene expression. This idea fits well with previous results from GWAS that 80% of the explained variation could be attributable to trait-associated variants located in regulatory regions [71]. In total, our DMR results provide a list of candidate genes to be further tested, especially those found in selective sweeps and interacting regions. To tease apart real DMR-phenotype associations from false, future efforts should focus on genotyping the methylation status of such loci across mapping populations while modeling SNP and DMR associations with phenotypes jointly.

In addition to our genome-wide approaches that identify a large number of novel DMRs, we also conducted functional validation at two well-studied candidate loci: *vgt1* and *tb1.* In both cases, our evidence showed that methylation affects physical interactions between the gene and intergenic regulatory regions. In particular, the maize alleles having low methylation levels exhibit interactive loops and increased expression of the downstream gene compared to highly methylated alleles in teosinte.

Collectively, our results suggest a meaningful functional role for methylation variation in maize. Genome-wide variation in methylation shows signs of weak natural selection and regions exhibiting variation explain considerable phenotypic variation. We also identify a large number of DMRs, many of which overlap with signals of selection during maize domestication and improvement as well as regions of the genome important for chromatin interaction. These results suggest that further investigation of the role of methylation in affecting genome-wide patterns of chromatin interaction and gene regulation is warranted, and that naturally occurring DMRs may provide a useful source of regulatory variation for crop improvement.

## METHODS

### Plant materials and DNA sequencing

We obtained a set of geographically widespread open pollinated landraces across Mexico (*N* = 17) from Germplasm Resources Information Network (GRIN) (**Table S1)**. The teosinte (Zea *mays* ssp. *parviglumis; N =* 20) were collected near Palmar Chico, Mexico [30]. We harvested the third leaf of the teosintes and Mexican landraces at the third leaf stage for DNA extraction using a modified CTAB procedure [72]. The extracted DNA was then sent out for whole genome sequencing (WGS) and whole genome bisulfite sequencing (WGBS) using Illumina HiSeq platform. Additionally, we obtained WGBS data for 14 modern maize inbred lines [6] and WGS data for the same 14 lines from the maize HapMap3 project [31].

### Sequencing data analysis

The average coverage for the WGS of the 20 teosintes and 17 landraces lines was about 20 ×. For these WGS data, we first mapped the cleaned reads to the B73 reference genome (AGPv4) [73] using BWA-mem [74] with default parameters, and kept only uniquely mapped reads. Then we removed the duplicated reads using Picard tools [75]. We conducted SNP calling using Genome Analysis Toolkit’s (GATK, version 4.1) HaplotypeCaller [76], in which the following parameters were applied: QD < 2.0, FS > 60.0, MQ < 40.0, MQRankSum < −12.5, and ReadPosRankSum < −8.0.

In order to improve the WGBS mapping rate and decrease the mapping bias, we replaced the B73 reference genome with filtered SNP variants using an in-house developed software — pseudoRef (https://github.com/yangjl/pseudoRef). Subsequently, we mapped reads to each corrected pseudo-reference genome using Bowtie2 [77] and kept only unique mapped reads. After filtering the duplicate reads, we extracted methylated cytosines using the Bismark methylation extractor and only retained sites with more than three mapped reads. The weighted methylation level was determined following the previously reported method [78].

### Population epigenetics modeling

Spontaneous epimutation changes (i.e. gain or loss of cytosine methylation) exhibit higher rate than genomic mutation [21, 69]. The standard population genetic methods designed for SNPs are thus inappropriate for population epigenetic studies. Here, we applied the analytical framework for hypermutable polymorphisms developed by Charlesworth and Jain [23]. Under this framework, the probability density of the methylated alleles was modeled as:

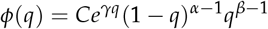

where *α* = 4*N_e_μ*, *β* = 4*N_e_v*, *γ* = 2*N_e_s*. *N_e_*, effective population size; *q*, frequency of the hypermethylation alleles; *μ*, forward epimutation rate (methylation gain); *v*, backward epimutation rate (methylation loss); *s*, selection coefficient. The constant *C* is required so that 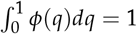.

We defined 100-bp tiles as a DNA methylation locus. To define the methylation status, we assumed that the methylation levels in a heterozygote individual falling into three mixture distributions (unmethylated, methylated, and heterozygote distributions). We employed an R add-on package “mixtools” and fitted the “normalmixEM” procedure to estimate model parameters [38]. Based on the converged results of the iterative expectation maximization algorithm (using the “normalmixEM” function), we decided to use 0.7 and 0.3 thresholds for heterozygote individuals (i.e., average methylation value> 0.7 for a 100-bp tile was determined as a methylated call and coded as 2; < 0.3 was determined as an unmethylated call and coded as 0; otherwise coded as 1). We also tested different cutoffs and found that the final methylation site frequency spectrum (mSFS) was insensitive to the cutoffs used. Similarly, we assumed two mixture distributions for inbred lines and used cutoff = 0.5 to determine methylated (coded as 1) and unmethylated (coded as 0) calls. With these cutoffs, we then constructed mSFS on genome-wide methylation loci. We also constructed interspecific (i.e., across maize, landrace, and teosinte populations) and intraspecific (i.e., within maize, landrace, and teosinte populations) mSFS.

To estimate three critical population epigenetic parameters (*μ*, *v*, and *s*) from observed mSFS, we implemented a Markov Chain Monte Carlo (MCMC) method (https://rpubs.com/rossibarra/mcmcbc). In the analyses, we selected a set of *N_e_ =* 50,000, 100,000, 500,000, and 1,000,000 [39, 40, 79, 80]. To test the prior values on the posterior distributions, we sampled *μ*, *v*, and *s* from exponential proposal distributions with different prior values of 10^2^, 10^4^, 10^5^, 10^8^, and 10^10^ (**Figure S8A** and lambda values of the scaled proposal distribution of 0.01, 0.05, and 0.1 (**Figure S8B**). We ran the model using a chain length of *N =* 1,000,000 iterations with the first 20% as burnin.

### Genome scanning to detect selective signals

We called SNPs using our WGS data and performed genome scanning for selective signals using XP-CLR method [81]. In the XP-CLR analysis, we used a 50-kb sliding window and a 5-kb step size. To ensure comparability of the composite likelihood score in each window, we fixed the number assayed in each window to 200 SNPs. We evaluated evidence for selections across the genome in three contrasts: teosinte vs landrace, landrace vs modern maize, and teosinte vs modern maize. We merged nearby windows falling into the 10% tails into the same window. After window merging, we considered the 0.5% outliers as the targets of selection.

We calculated *F_ST_* using WGS data using VCFtools [82]. In the analysis, we used a 50-kb sliding window and a 5-kb step size.

### DMR detection and GO term analysis

We used a software package ‘metilene’ for DMR detection between two populations [44]. To call a DMR, we required it contained at least eight cytosine sites with < 300-bp in distance between two adjacent cytosine sites, and the average of methylation differences between two populations should be > 0.4 for CG and CHG sites. Finally, we required a corrected *P*-value < 0.01 as the cutoff.

We conducted gene ontology (GO) term analysis on selected gene lists using AgriGO2.0 with default parameters [83]. We used the significance cutoff at *P*-value < 0.01.

### Linkage disequilibrium (LD) analysis between DMR and local sNPs

To test the relationship between DMRs and selective sweeps, we conducted LD analysis using SNPs located 1 kb upstream and downstream of each DMR. A DMR was determined as in LD if there are at least three SNPs displayed significant correlations with this DMR (permutation P-value < 0.0l).

### HichiP sequencing library construction

We constructed the teosinte HiChIP library according to the protocol developed by Mumbach *et al.* [84] with some modifications. The samples we used were two weeks aerial tissues collected from a teosinte accession (Ames 21809) that were planted in the growth chamber under the long-day condition (15h day time and 9h night time) at the temperature (25°C at day time and 20°C at night time). After tissue collection, we immediately cross-linked it in a 1.5 mM EGS solution (Thermo, 21565) for 20 min in a vacuum, followed by 10 min vacuum infiltration using 1% formaldehyde (Merck, F8775-500ML). To quench the EGS and formaldehyde, we added a final concentration of 150 mM glycine (Merck, V900144) and infiltrated by vacuum for 5 min. Then, cross-linked samples were washed five times in double-distilled water and flash-frozen in liquid nitrogen.

To isolate the nuclear from cross-linked tissues, we used the methods as described previously [33]. After obtaining the purified nuclear, we resuspended it in 0.5% SDS and used 10% Triton X-100 to quench it, and then performed digestion, incorporation, and proximity ligation reactions as previously described [84]. We used two antibodies H3K4me3 (Abcam, ab8580) and H3K27ac (Abcam, ab4729) to pull down the DNA. And then, we purified DNA with the MinElute PCR Purification Kit (QIAGEN, Cat No. 28006) and measured the DNA concentration using Qubit. To fragment and capture interactive loops, we used the Tn5 transposase kit (Vazyme, TD501) to construct the library. We then sent the qualified DNA libraries for sequencing using the Illumina platform.

### ChlP-seq and HiChIP data analysis

We obtained ChIP-seq data from the B73 shoot tissue [33] and then aligned the raw reads to B73 reference genome (AGPv4) using Bowtie2 [85]. After alignment, we removed the duplicated reads and kept only the uniquely mapped reads. By using the uniquely mapped reads, we calculated read coverages using deepTools [86].

For the teosinte HiChIP sequencing data, we first aligned the raw reads to the B73 reference genome (AGPv4) using HiC-Pro [87], and then processed the valid read pairs to call interactive loops using hichipper pipeline [88] with a 5-kb bin size. After the analysis, we filtered out the non-valid loops with genomic distance less than 5 kb or larger than 2 Mb. By using the mango pipeline [89], we determined the remaining loops with three read pairs supports and the FDR < 0.01 as the significant interactive loops.

### 4c-seq library construction and data analysis

To validate the physical interaction between tb1-DMR and *tb1* gene, we performed 4C-seq experiments using landrace samples. We constructed the 4C-seq libraries using restriction enzymes of NlaIII and *DpnII.* The primer sequences for the *tb1* bait region were the same as previously described [33]. After sequencing, we aligned the reads to the B73 reference genome and then processed the uniquely mapped reads using 4C-ker program [90].

### Kinship matrices and variance components analysis

We estimated the variance components explained by SNP sets residing in DMRs using the maize Nested Association Mapping (NAM) population [91, 92]. We downloaded the phenotypic data (/iplant/home/glaubitz/RareAlleles/genomeAnnos/VCAP/phenotypes/ NAM/familyCorrected), consisting of Best Linear Unbiased Predictors (BLUPs) for different traits ([42]), and imputed genotypic data (/iplant/home/glaubitz/RareAlleles/genomeAnnos/VCAP/genotypes/NAM/namrils_projected_hmp31_MAF02mnCnt2500.hmp.txt.gz) [31] from CyVerse database as described in Panzea (www.panzea.org).

In the analysis, we mapped SNPs to the invariable hypermethylated, invariable hypomethylated, and variable methylated regions. For each SNP set, we calculated an additive kinship matrix using the variance component annotation pipeline implemented in TASSEL5 [93]. We then fed these kinship matrices along with the NAM phenotypic data to estimate the variance components explained by SNP sets using a Residual Maximum Likelihood (REML) method implemented in LDAK [43].

### Dual-luciferase transient expression assay in maize protoplasts

To investigate the effect of DMRs on gene expression, we performed a dual-luciferase transient expression assay in maize protoplasts. We used the pGreen II 0800-LUC vector [94] for the transient expression assay with minor modification, where a minimal promoter from cauliflower mosaic virus (mpCaMV) was inserted into the upstream of luciferase (LUC) to drive LUC gene transcription. In the construct, we employed the *Renillia luciferase (REN*) gene under the control of 35S promoter from cauliflower mosaic virus (CaMV) as an internal control to evaluate the efficiency of maize protoplasts transformation. We amplified the selected DMR sequences from B73 and then inserted them into the control vector at the restriction sites *KpnI/XhoI* upstream of the mpCaMV, generating the reporter constructs.

We planted B73 in the growth chamber and kept the plants in the darkness at the temperature of about 20°C (night) and 25°C (day) to generate etiolated plants. Protoplasts were isolated from the 14-day-old leaves of B73 etiolated seedlings following the protocol [95]. Subsequently, we transformed 15 ug plasmids into the 100 ul isolated protoplasts using polyethylene glycol (PEG) mediated transformation method [95]. After 16 hours infiltration, we measured the LUC and REN activities using dual-luciferase reporter assay reagents (Promega, USA) and a GloMax 20/20 luminometer (Promega, USA). Finally, we calculated the ratios of LUC to REN. For each experiment, we included five biological replications.

### Experimental validation of the tb1-DMR

We performed Chop-PCR (DNA methylation-sensitive restriction endonuclease digestion followed by PCR) to validate DNA methylation at tb1-DMR locus in different tissues of modern maize inbred line W22 and teosinte 8759. We collected the leaf tissue at the third leaf stage and immature ears of ≈5 cm in length. To evaluate the methylation level of tb1-DMR locus, we treated 1 *μ*g purified genomic DNA using the EpiJETTM DNA Methylation Analysis Kit (*MspI/HpaII*) (Thermo Scientific, K1441) following manufacturer’s instructions. The primer sequences for PCR were ACACGCACGAAGGGTTACAG (forward) and CAGTGCTCCCTGGGTCAAA (reverse).

## Statistical analyses

All the statistical tests were performed using R software (V3.6.2, https://www.r-project.org/).

## CODE AVAILABILITY

The code used for analyses can be accessed through GitHub (https://github.com/jyanglab/msfs_teo).

## ACKNOWLEDGEMENTS

J.Y. is supported by the Agriculture and Food Research Initiative Grant number 2019-67013-29167 from the USDA National Institute of Food and Agriculture, the National Science Foundation under award number OIA-1557417 for Center for Root and Rhizobiome Innovation (CRRI), Institutional Development Award (IDeA) from the National Institute of General Medical Sciences of the National Institutes of Health under Grant number P20GM103476, and the University of Nebraska-Lincoln Start-up fund and the Layman seed award. J.R.-I. is supported by NSF grant 1546719 and USDA Hatch project CA-D-PLS-2066-H. This work was conducted using the Holland Computing Center of the University of Nebraska-Lincoln Start-up, which receives supports from the Nebraska Research Initiative. We would like to thank Mike May for help developing the MCMC approach used here, the helpful discussion in J.R.-I.’s REHAB, and constructive suggestions from anonymous reviewers.

## AUTHOR CONTRIBUTIONS

J.Y. and J.R.-I. designed this work. J.L., Q.L., N.M.S., and D.W. generated the data. H.L. and M.Z. produced the teosinte HiChIP libraries. G.X., J.R.-I., and J.Y. analyzed the data. N.M.S. provided conceptual advice. J.Y., G.X. and J.R.-I. wrote the manuscript.

## SUPPORTING INFORMATION

### SUPPORTING TABLES

**Table S1.** Teosinte, landrace, modern maize samples used in this study. (https://github.com/jyanglab/msfs_teo/blob/master/table/Table_S1_samples_for_sequencing.xlsx)

**Table S2.** Population-wide DMRs in the CG and CHG contexts. (https://github.com/jyanglab/msfs_teo/blob/master/table/Table_S2_DMR.xlsx)

**Table S3.** Selective sweeps detected between populations. (https:https://github.com/jyanglab/msfs_teo/blob/master/table/Table_S3_Selective_sweep.xlsx)

**Table S4.** Linkage disequilibrium (LD) analysis between DMR and local SNP. (https://github.com/jyanglab/msfs_teo/blob/master/table/Table_S4_DMR_LD.xlsx)

**Table S5.** The list of genes with CG teosinte-maize DMRs located at their exonic regions. (https://github.com/jyanglab/msfs_teo/blob/master/table/Table_S5_229_Exonic_DMR_Genes.xlsx)

**Table S6.** The list of genes exhibiting interactive loops between genes and hypomethylated DMRs in maize located at the intergenic regions. (https://github.com/jyanglab/msfs_teo/blob/master/table/Table_S6_60_Genes_interactive_with_intergenic_hypo_DMR.xlsx)

**Table S7.** Flowering time candidate genes located at sweep and interacting DMRs. (https://github.com/jyanglab/msfs_teo/blob/master/table/Table_S7_FT_candidate_gene_at_sweep_interaction_DMR.xlsx)

### SUPPORTING FIGURES

**Fig. S1.**
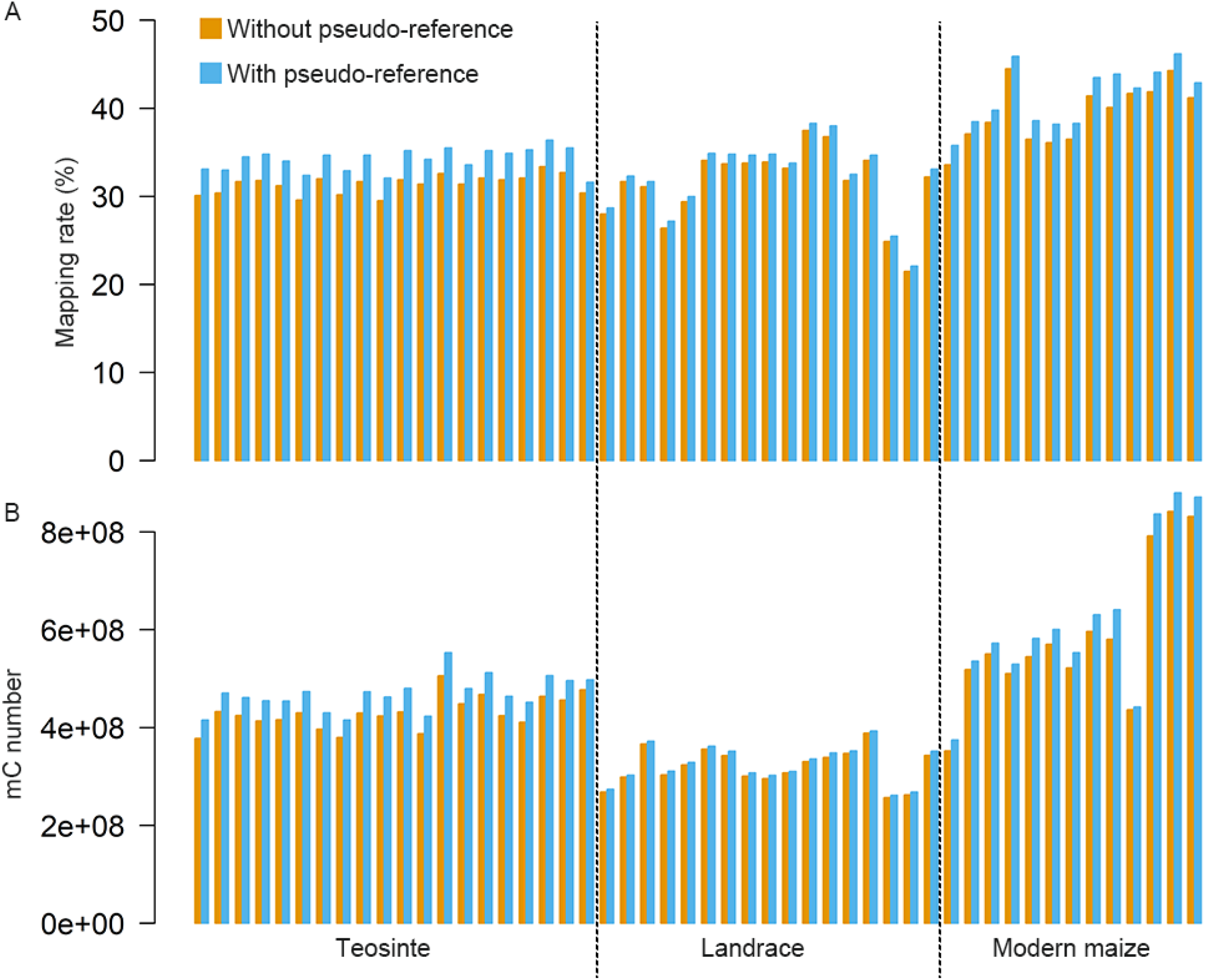
Comparison of mapping rates (A) and number of methylated cytosine (mC) sites (B) with and without using pseudoreference genome in different populations. B73 reference genome (AGPv4) was used in the analyses.

**Fig. S2.**
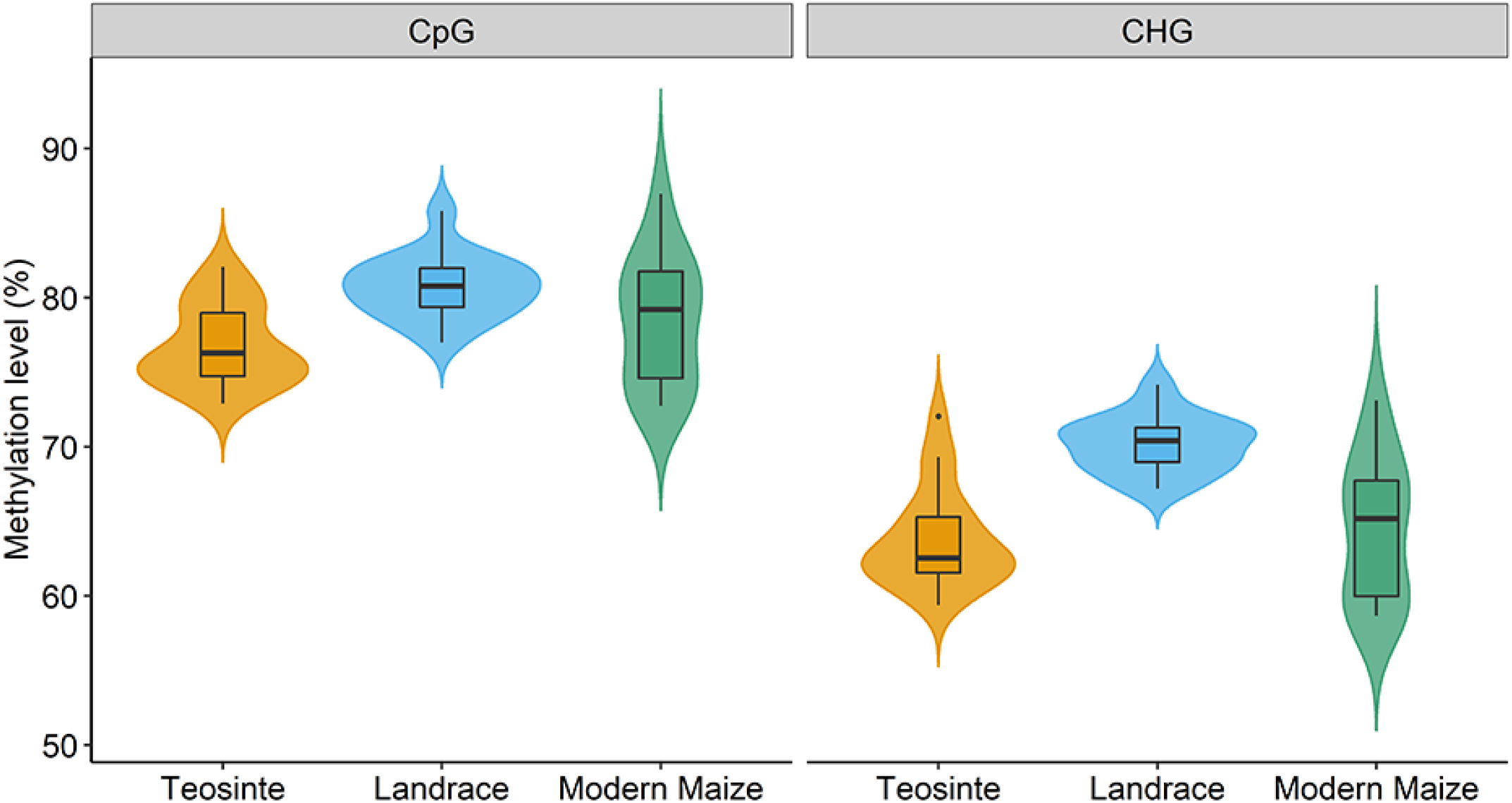
Distributions of levels of DNA methylation in teosinte, landrace, and modern maize populations. Left panel denotes results from CG sites and right panel denotes results from CHG sites.

**Fig. S3.**
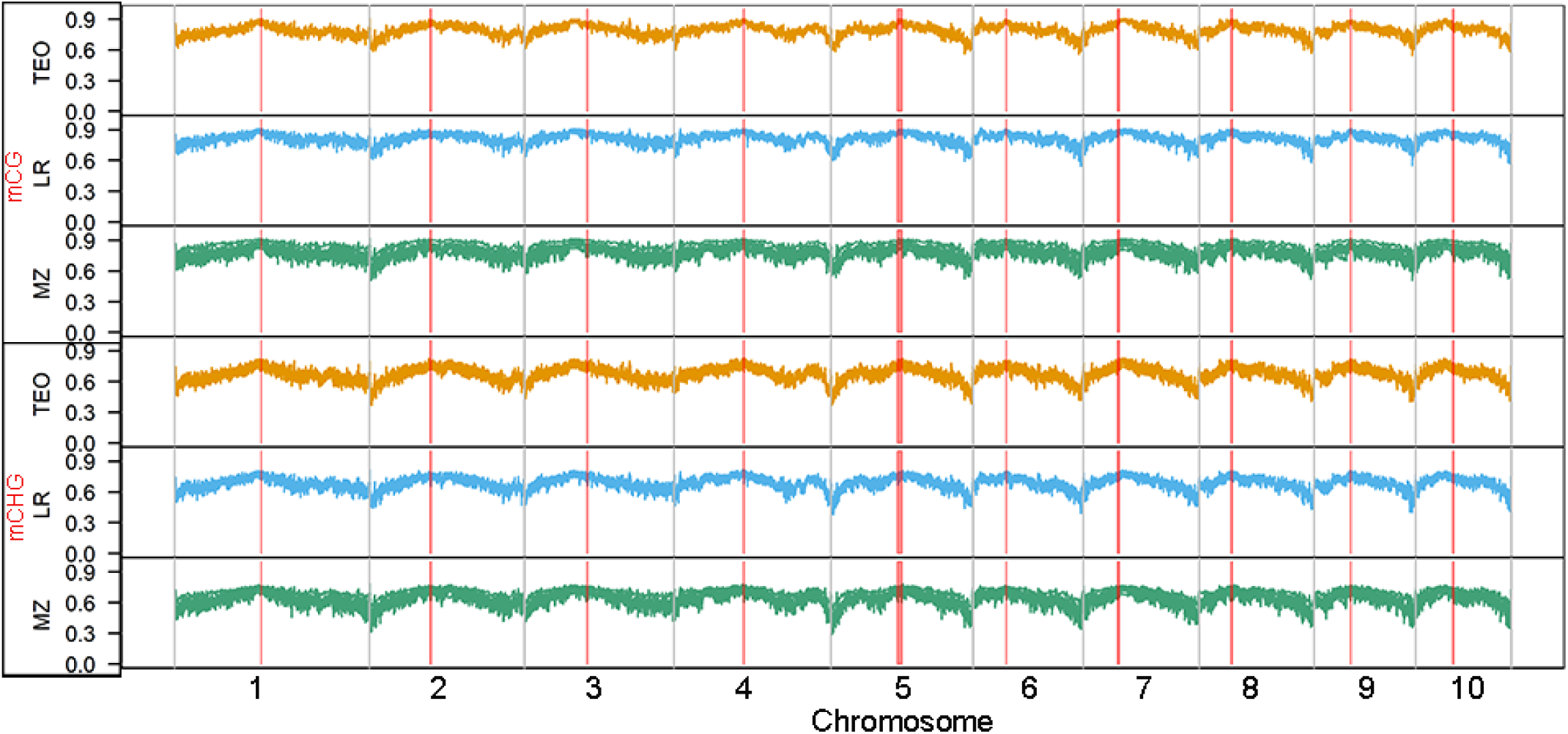
Genome-wide distributions of DNA methylation across 10 maize chromosomes. TEO, LR, and MZ represent teosinte, landrace and modern maize populations, respectively. Red vertical lines indicated the pericentromeric regions.

**Fig. S4.**
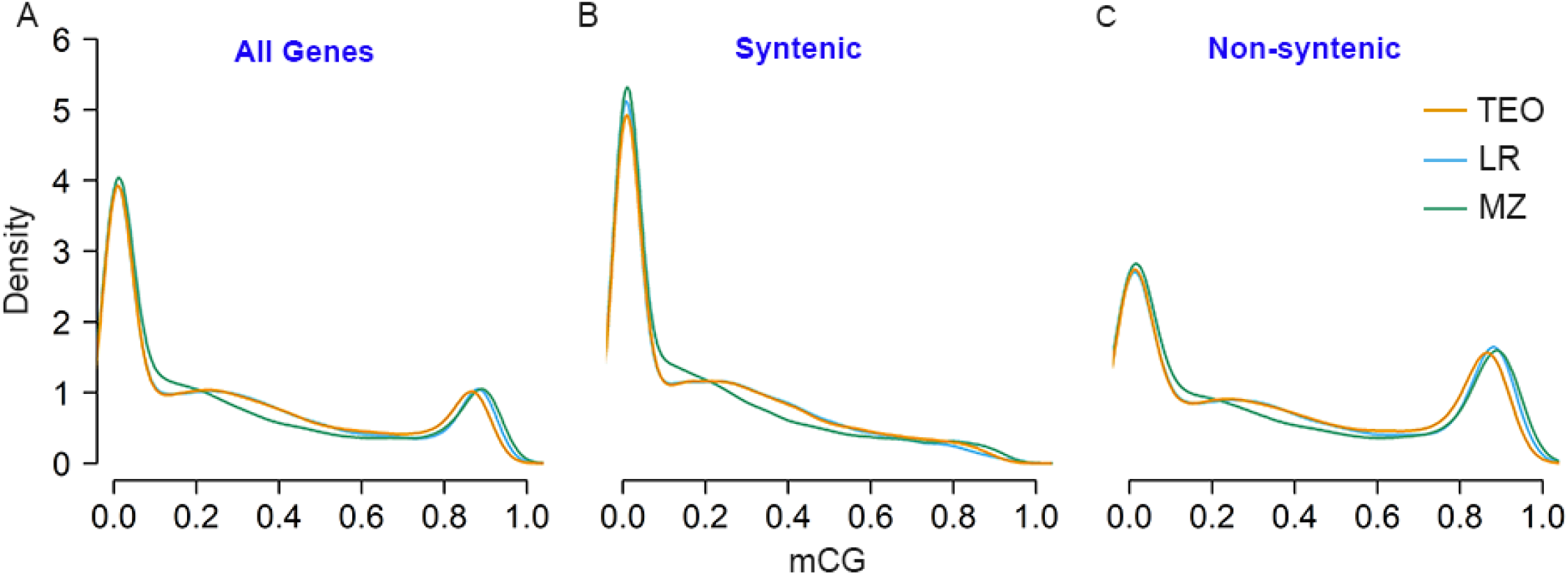
Density plots of CG methylation in gene body for all the annotated maize genes (A) and for syntenic (B) and non-syntenic genes (C). TEO, LR, and MZ represent teosinte, landrace and modern maize populations, respectively. The syntenic and nonsyntenic orthologs was determined by comparing maize with Sorghum.

**Fig. S5.**
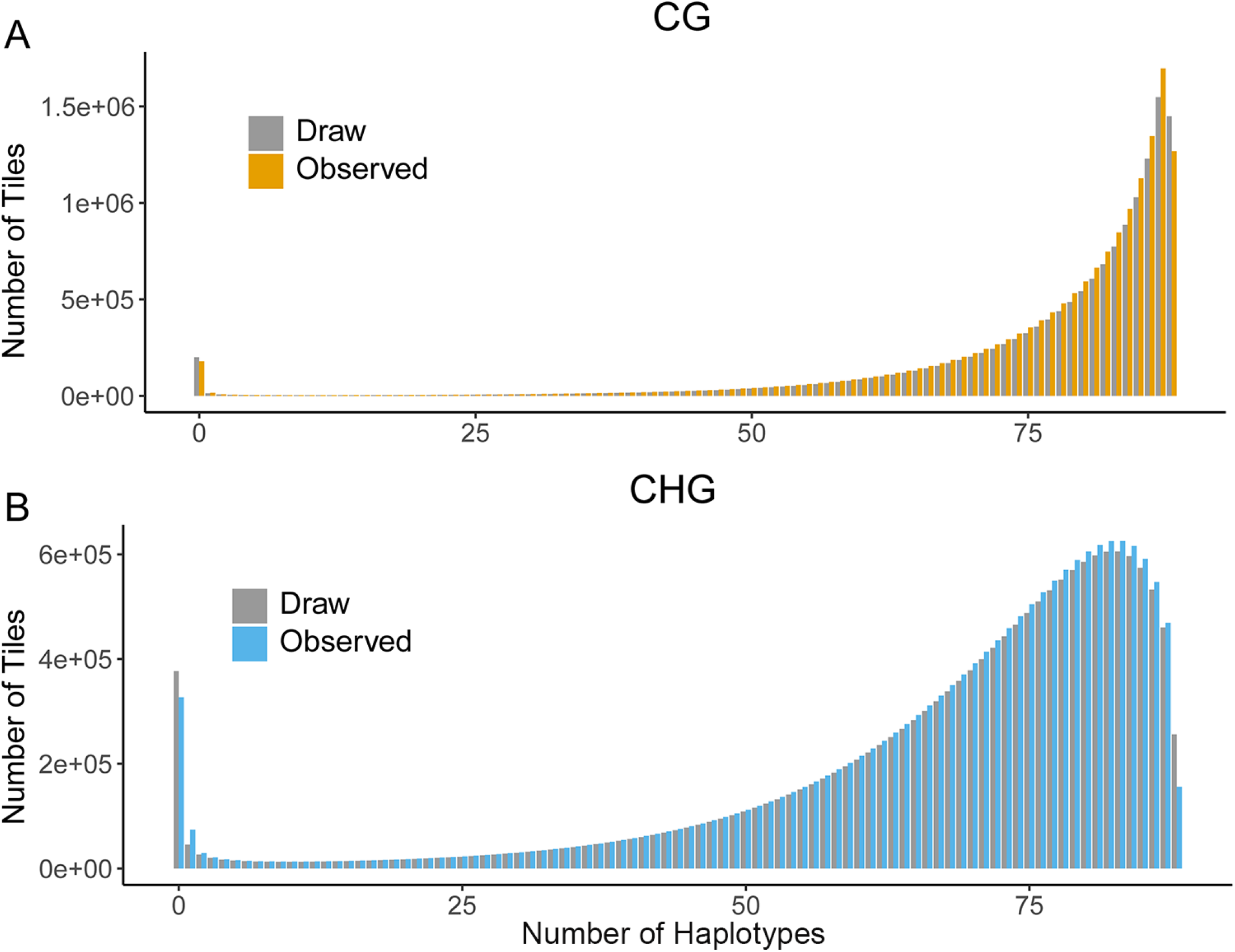
Observed and posterior methylome site frequency spectra. The posterior mSFS was calculated using parameters drawn from the 1,000,000th iteration (Ne=50,000).

**Fig. S6.**
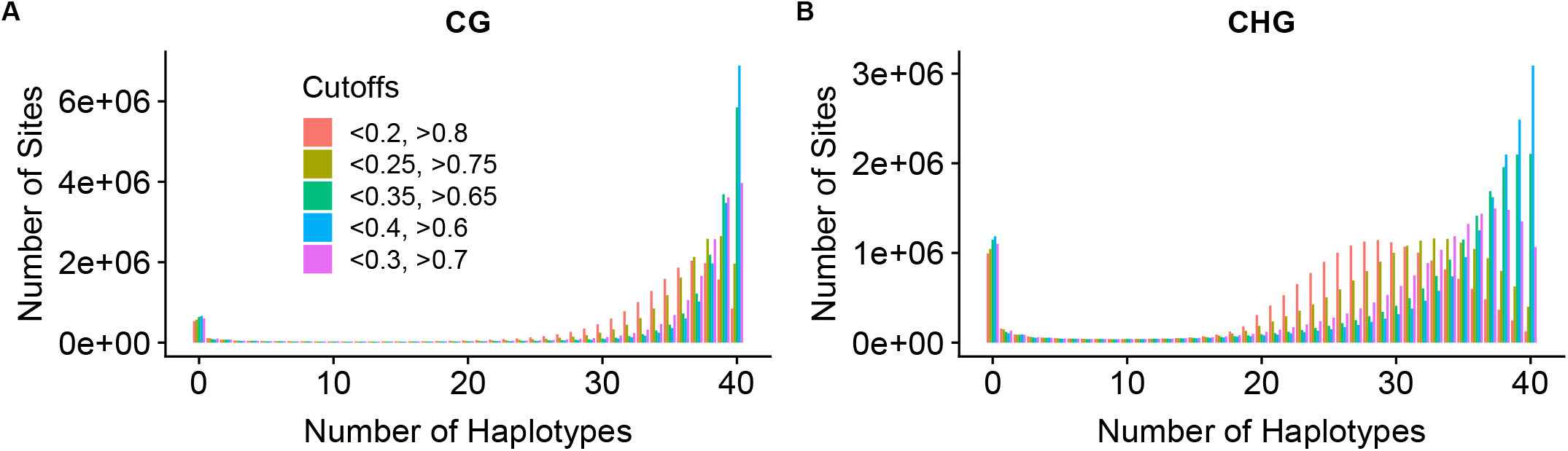
Sensitivity tests using different cutoffs. Distributions of mSFS using different thresholds to determine the methylated, unmethylated, and heterozygote for the 100-bp tiles under CG (**A**) and CHG (**B**) contexts.

**Fig. S7.**
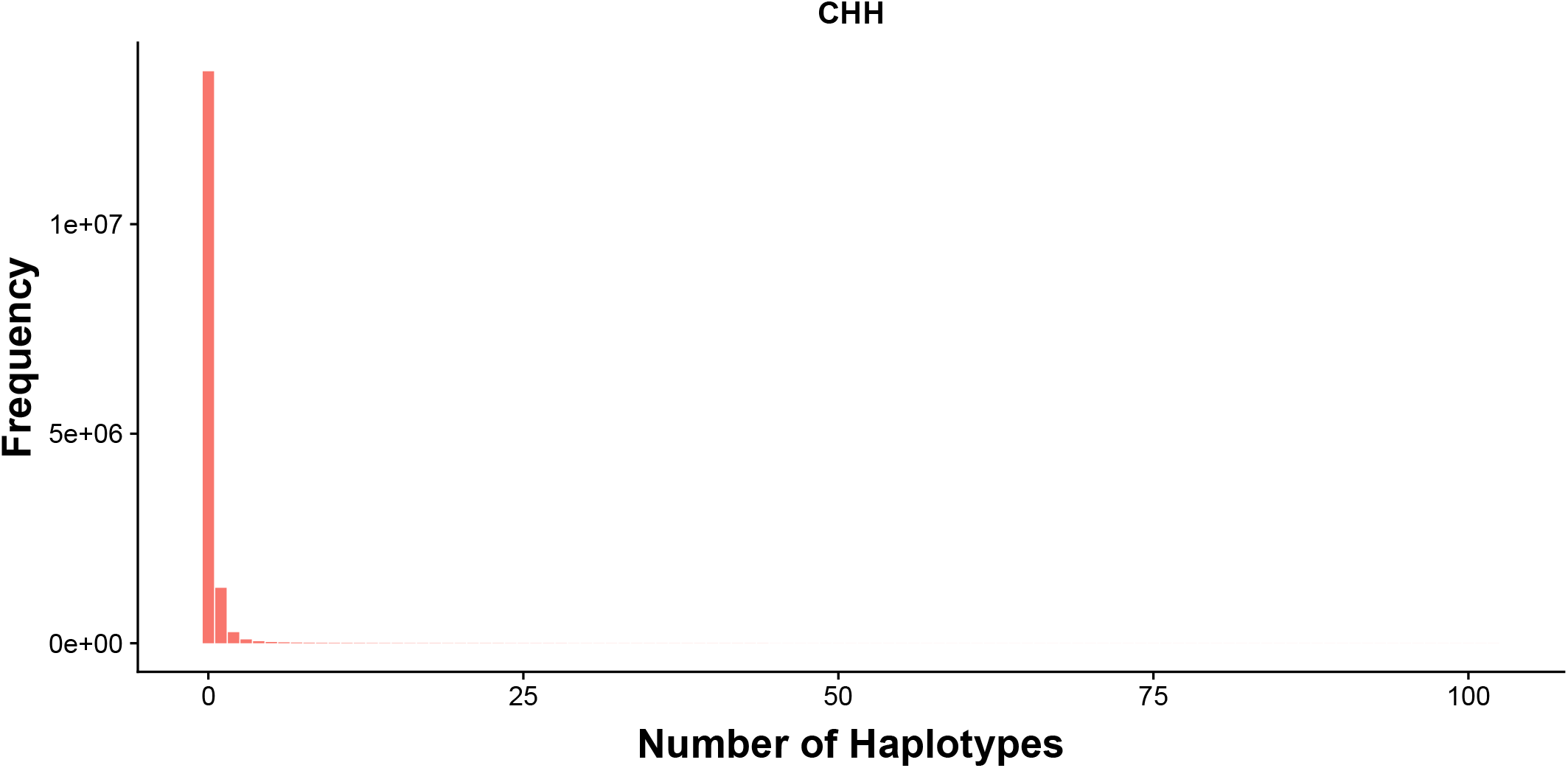
Methylome site frequency spectrum under the CHH context. The distribution is highly skewed towards the unmethylated status for CHH sites.

**Fig. S8.**
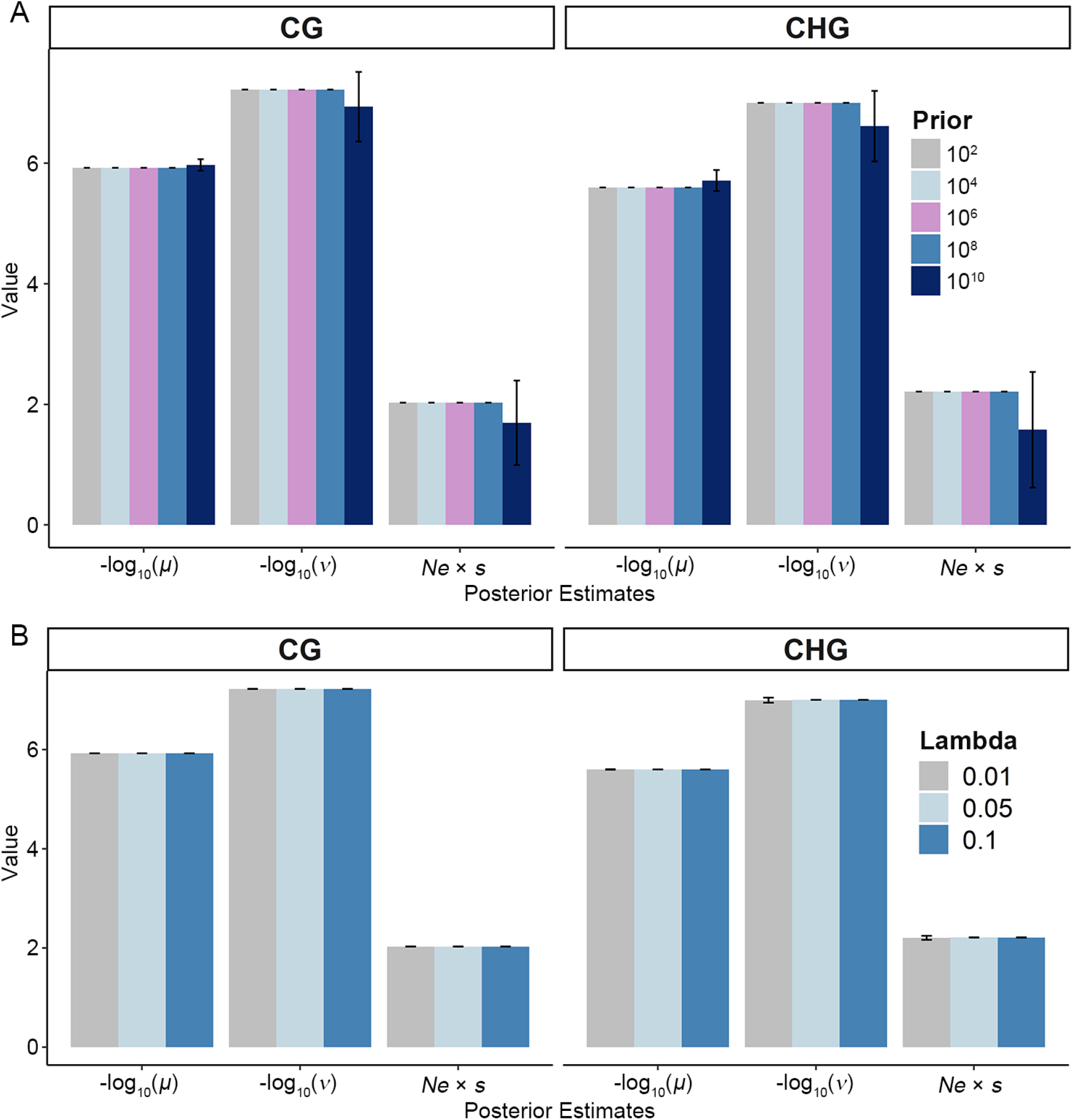
The effect of prior values on posterior parameter estimations. (**A**) Prior values of 10^2^, 10^4^, 10^5^, 10^8^, and 10^10^ were used for the exponential proposal distributions. (**B**) Lambda values of the scaled proposal distribution of 0.01, 0.05, and 0.1 were used. Error bars indicate standard deviations. Data available at (N = 1,600 for each bar).

**Fig. S9.**
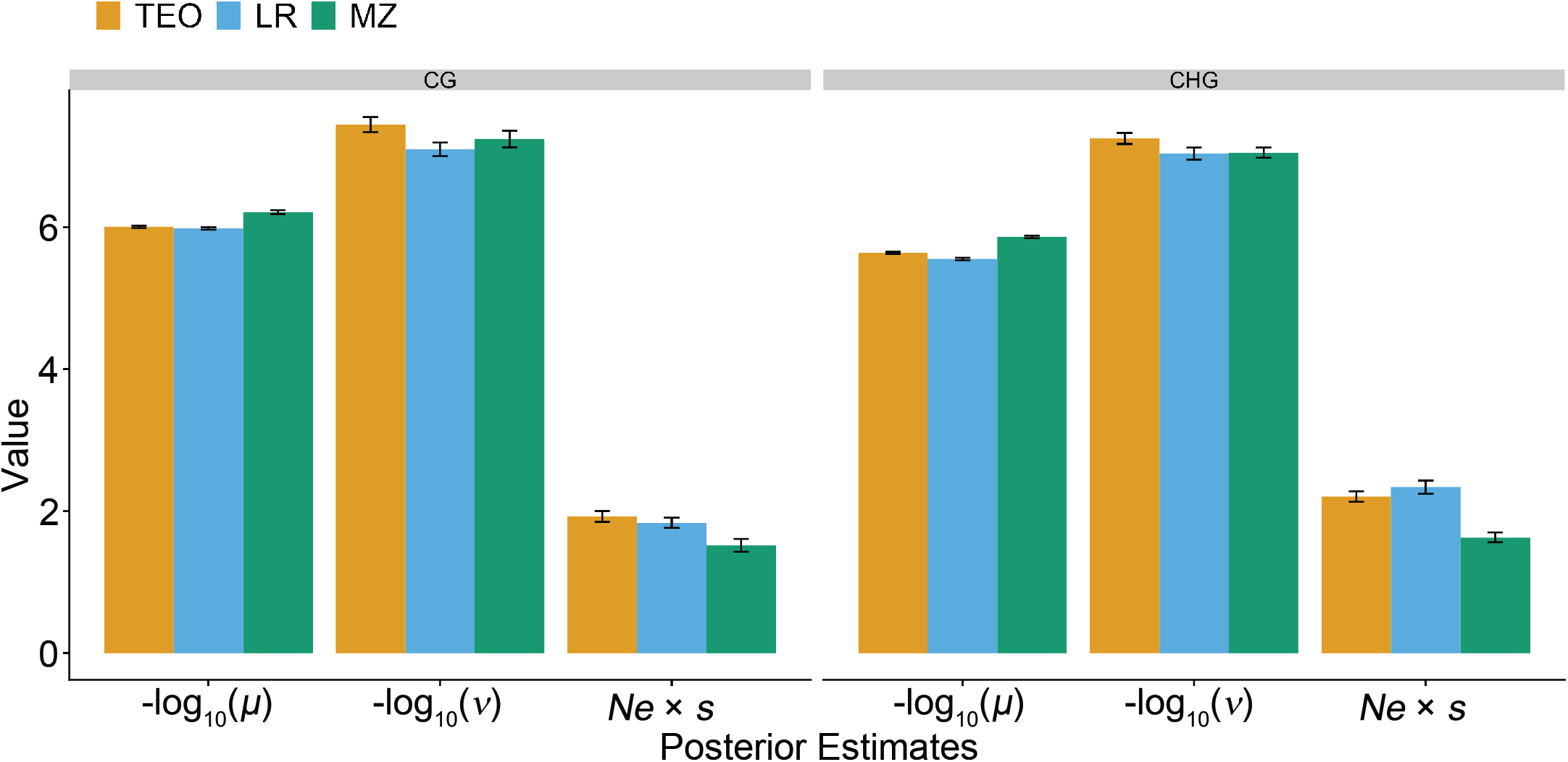
Population genetic parameter inference using each individual population. Posterior estimators of mean values and standard deviations for *μ, v,* and Ne × *s* for CG and CHG sites. Values were estimated using MCMC approach with 25% burnin. Error bars indicate standard deviations. Data available at (N = 1,600 for each bar).

**Fig. S10.**
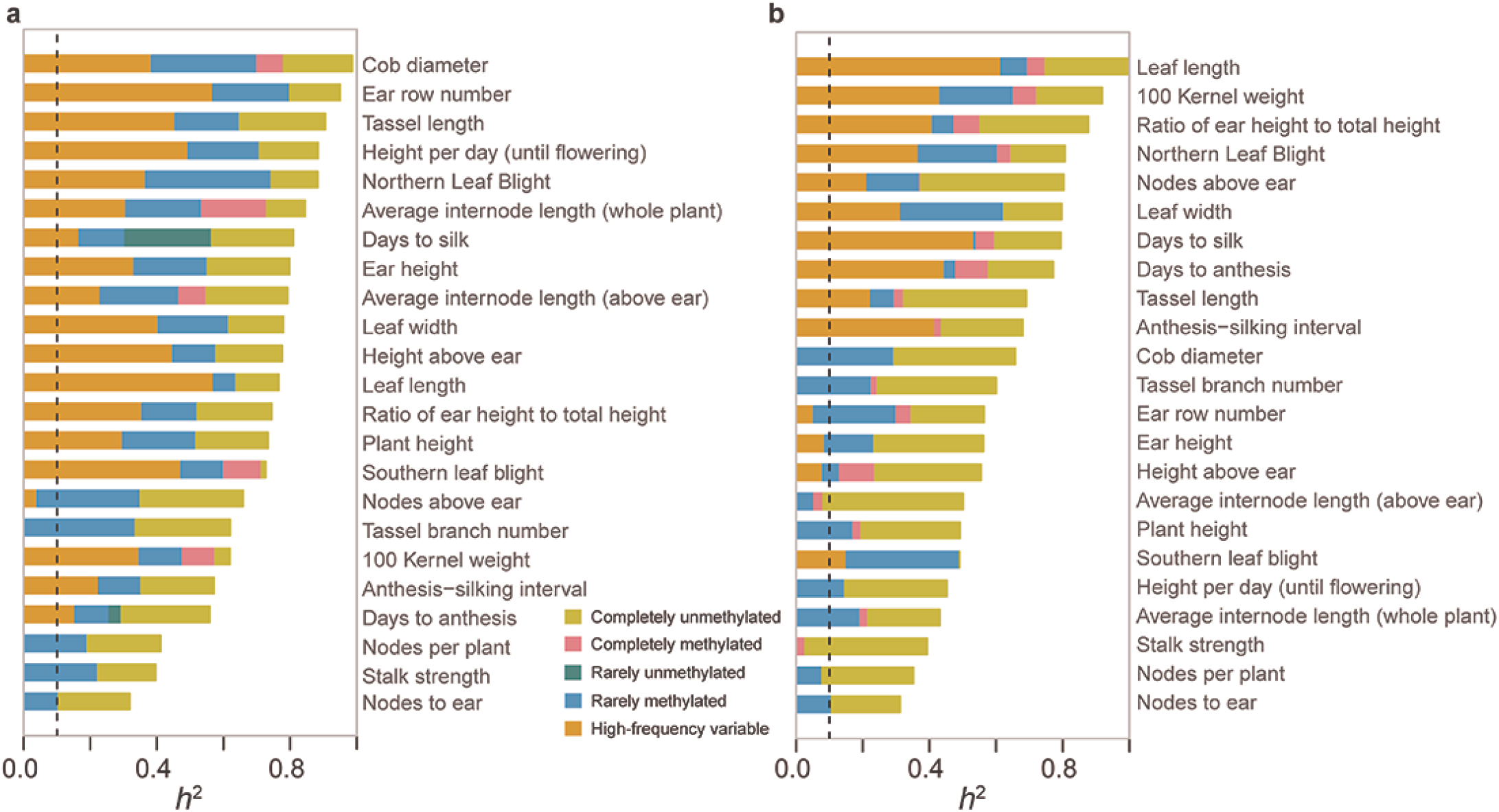
Proportion of genetic variances explained by SNP subsets residing in different genomic regions with different DNA methylation status. The proportion of genetic variance explained (h^2^) by different SNP subsets under CG (**A**) and CHG (**B**) contexts.

**Fig. S11.**
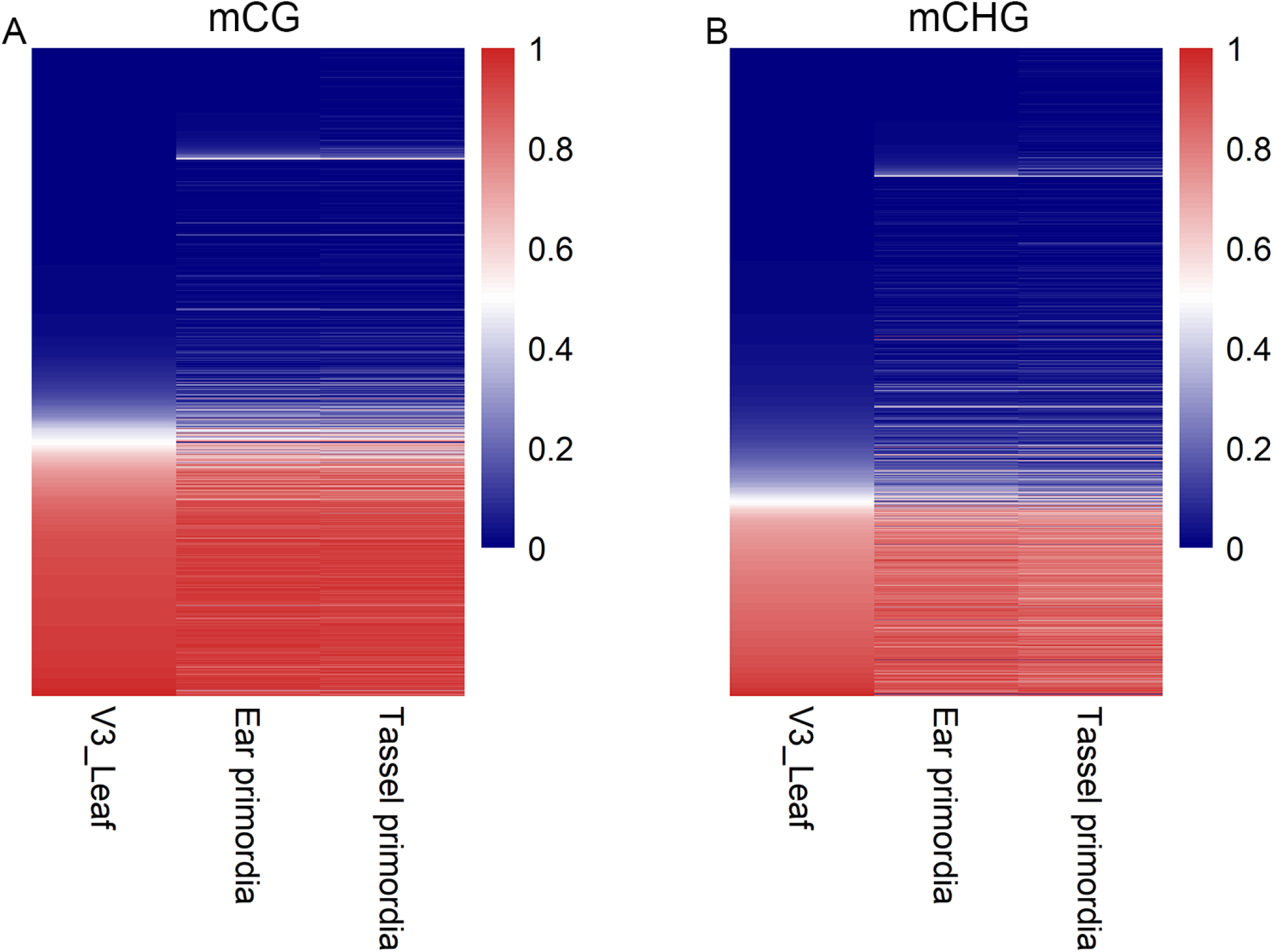
The methylation patterns of the DMRs across different tissues. DNA methylation levels in CG (**A**) and CHG (**B**) of each DMR across three tissues. The colors in the heatmap indicate the high (red) or low (blue) DNA methylation levels.

**Fig. S12.**
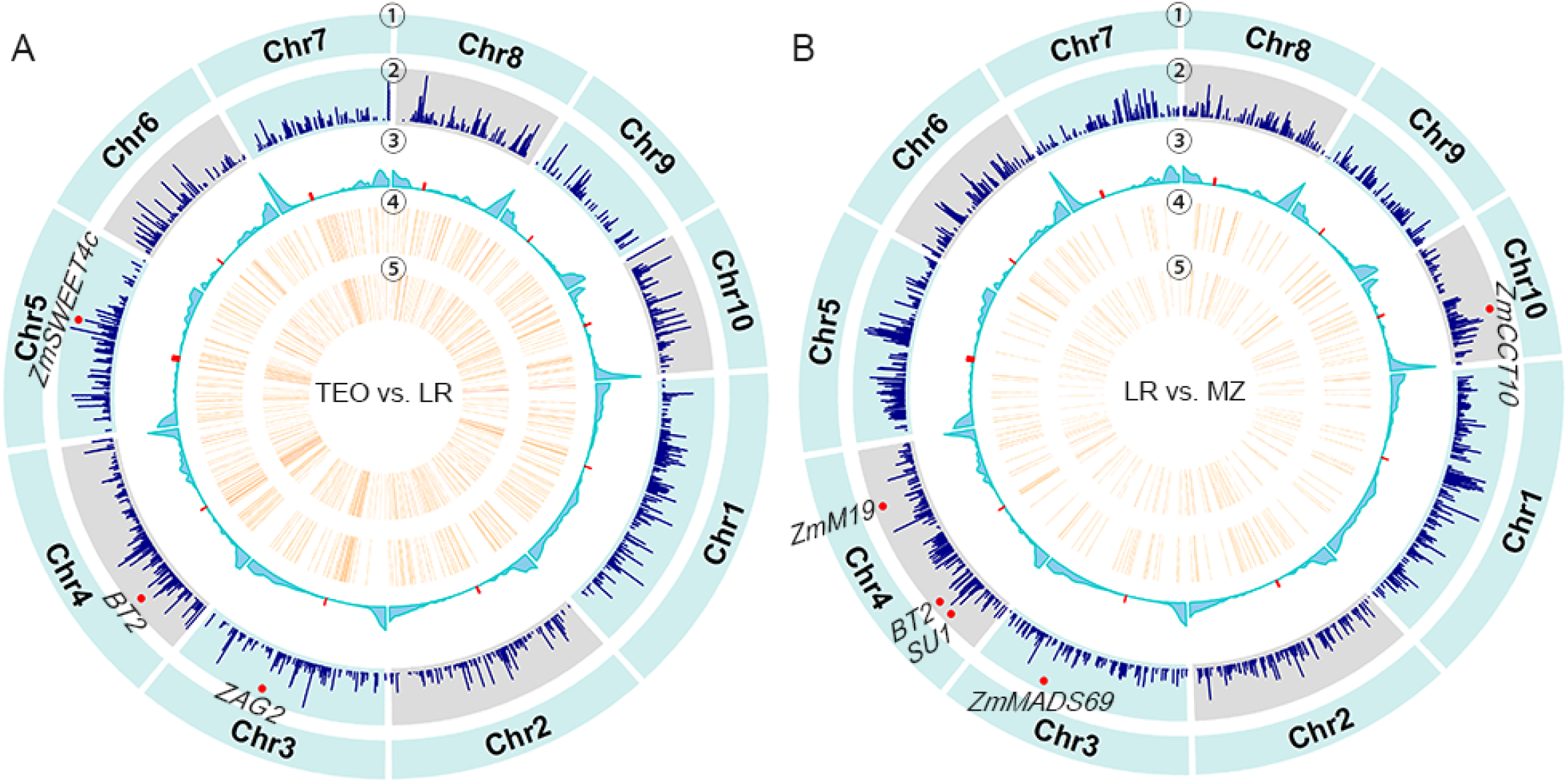
Landscape of selection signals and DNA methylation variation across maize genome. Genome-wide distributions of selective sweeps, DMRs across ten maize chromosomes detected by teosinte vs. landrace (**A**) and landrace vs. modern maize (**B**). From outer to inner circles: ① chromosome names, ② selective sweeps, ③ recombination rate, and the density of DMRs (number per 1-Mb) in ④ CG and ⑤ CHG contexts. Red dots at the second track indicated the physical positions of the known genes located within the selective sweeps. Red dots at the third track indicated the centromeric regions.

**Fig. S13.**
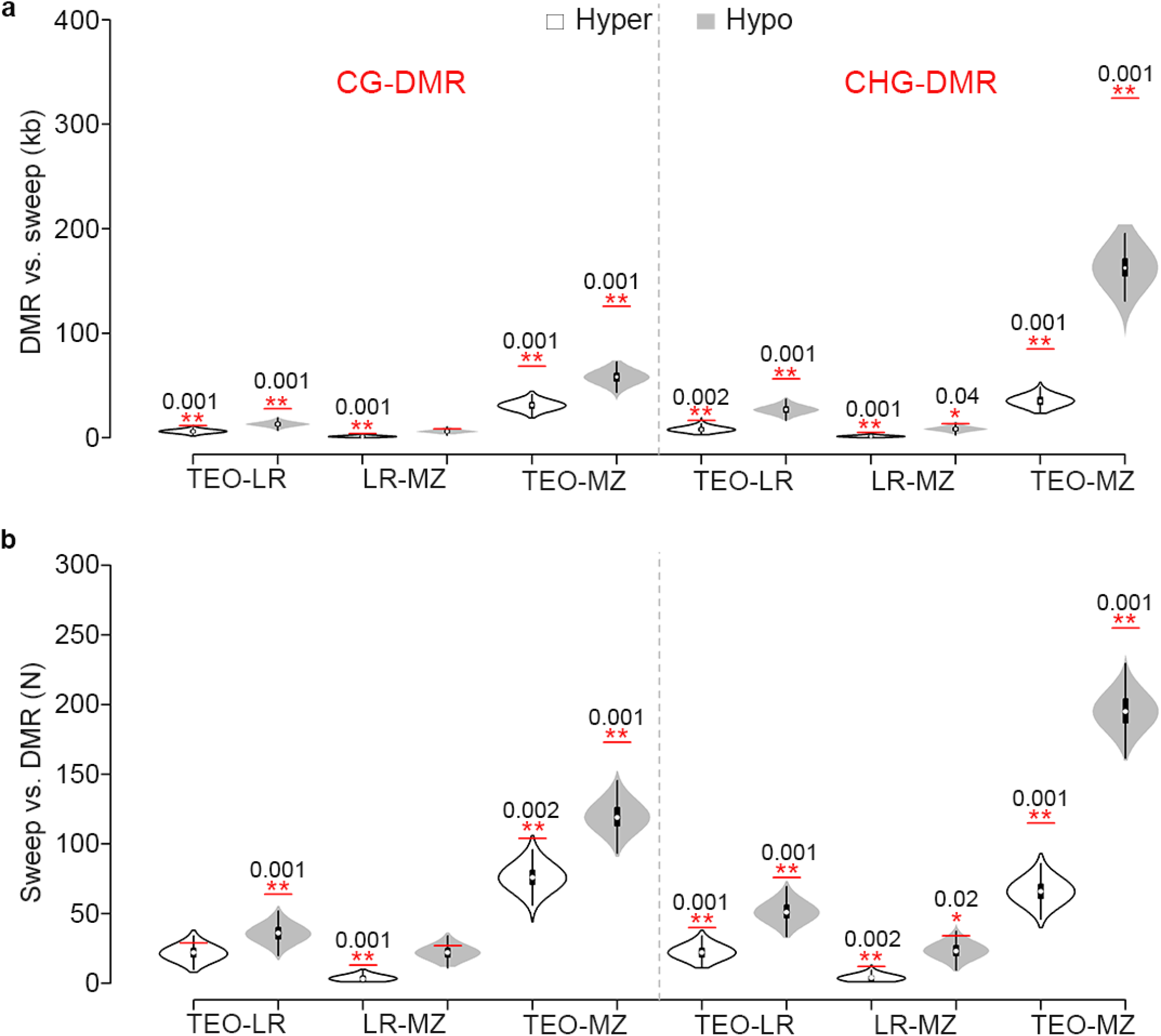
Comparison between DMRs and selective sweeps. (**A**) The overlapped base-pairs between DMRs and selective sweeps. (**B**) The number of sweeps that overlapped with DMRs. Red horizontal bars indicated the observed values and violin plots showed the 1,000 one-sided permutation results using randomly selected mappable regions from the genome. Red asterisks indicated the statistical significance with one asterisk denoting P-value < 0.05 and two asterisks denoting P-value < 0.01. The numbers above the asterisks indicate the exact P-values. Hyper- and hypomethylation were defined based on maize.

**Fig. S14.**
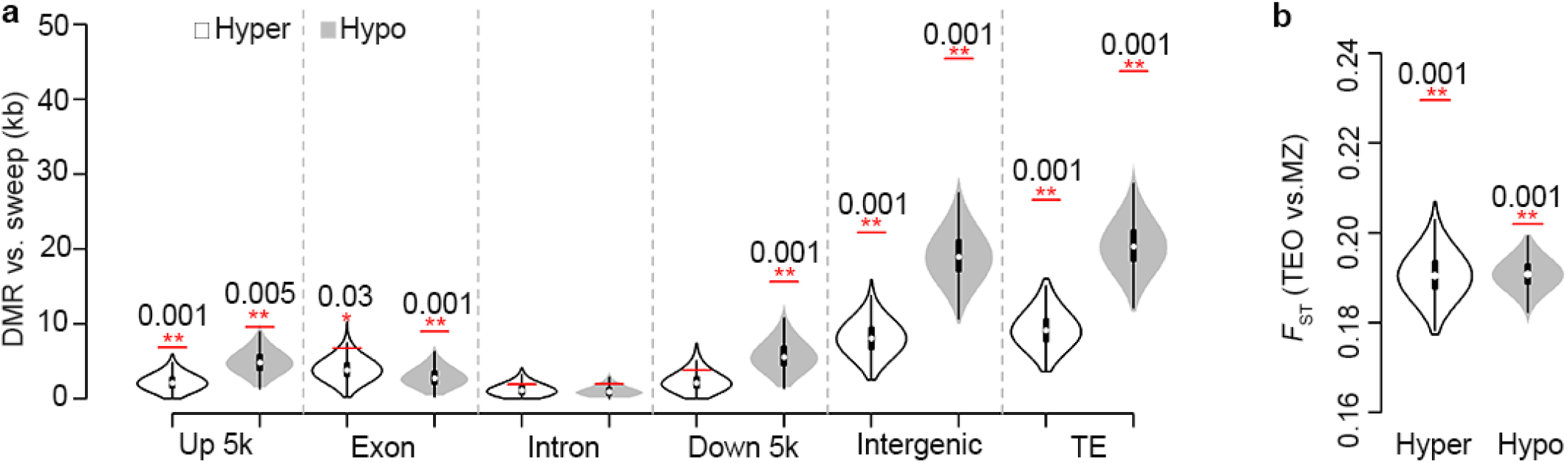
Selection on differentially methylated regions. (**A**) Overlaps between teosinte-maize DMRs and selective sweeps breaking down into different genomic features. (**B**) Mean *F_ST_* values of teosinte-maize DMRs that were hyper- and hypomethylated in maize. Red horizontal bars indicated the observed values and violin plots showed the 1,000 one-sided permutation results using randomly selected regions sharing the similar genomic features as the DMRs. Red asterisks indicated the statistical significance with one asterisk denoting P-value < 0.05 and two asterisks denoting P-value < 0.01. The numbers above the asterisks indicate the exact P-values.

**Fig. S15.**
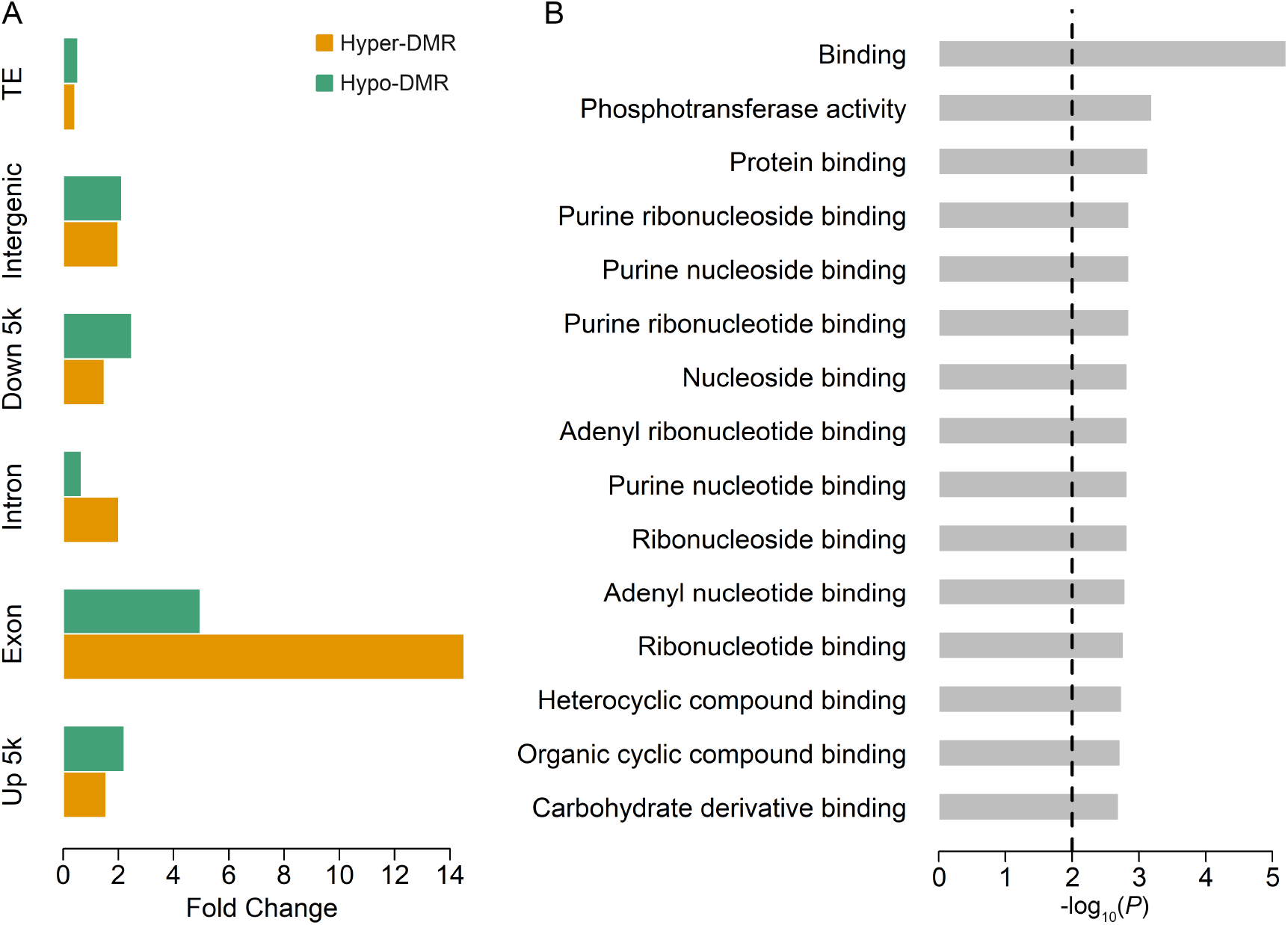
Teosinte-maize CG DMRs and their associated functional features. (**A**) Fold changes of mappable DMR length relative to the mean values from 1,000 permutations. (**B**) Result of GO term enrichment test using genes exhibiting an exonic DMR. Vertical dashed line indicates the significance cutoff (Fisher’s exact test, P-value=0.01).

**Fig. S16.**
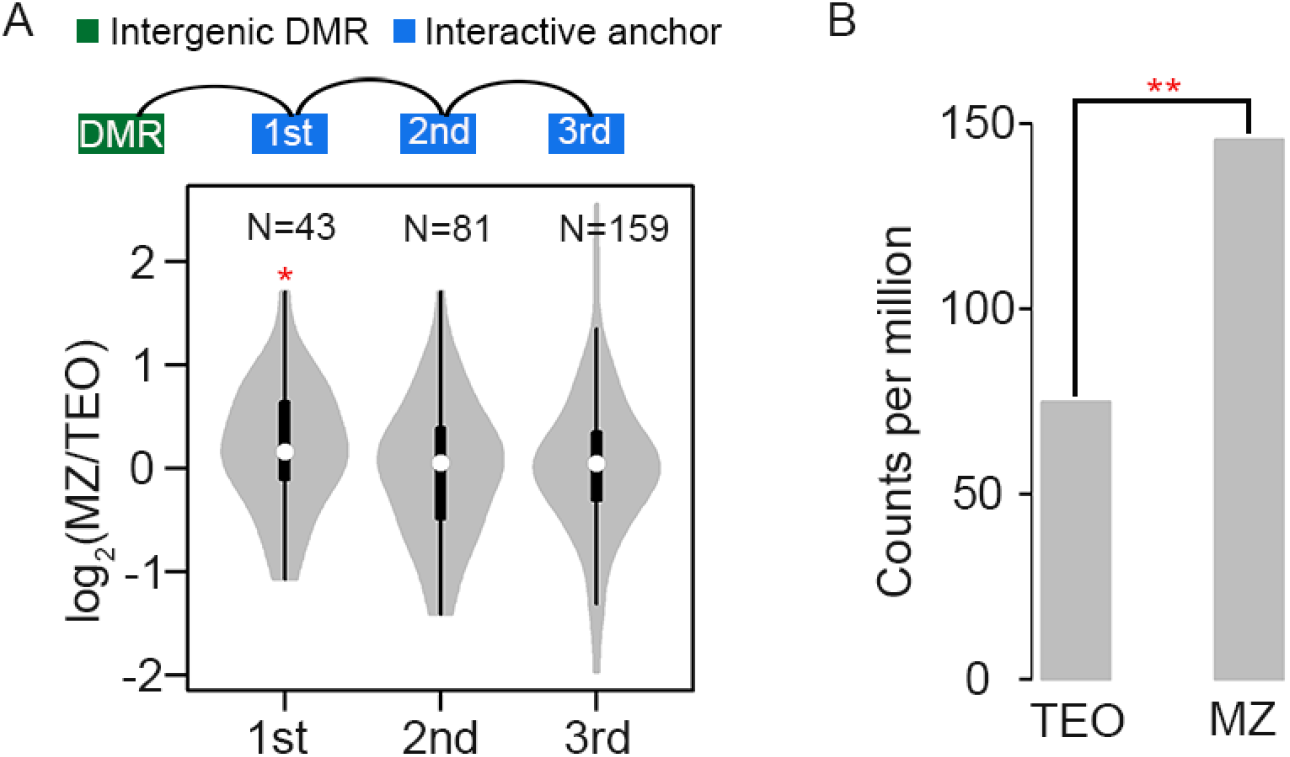
Intergenic CG DMR altered downstream gene expression. (**A**) Contrast of the gene expression levels in maize relative to teosinte. In the upper panel, the schematic diagram shows the genes that involved in the 1st, 2nd, and 3rd level interactions with maize hypomethylated DMRs located in intergenic regions. Red asterisk indicates the statistically significance (two-sided paired t-test: P-value < 0.05). (**B**) Gene expression level of *Zm00001d018036* in teosinte and modern maize (Binomial test, P-value = 4.6e^−141^).

**Fig. S17.**
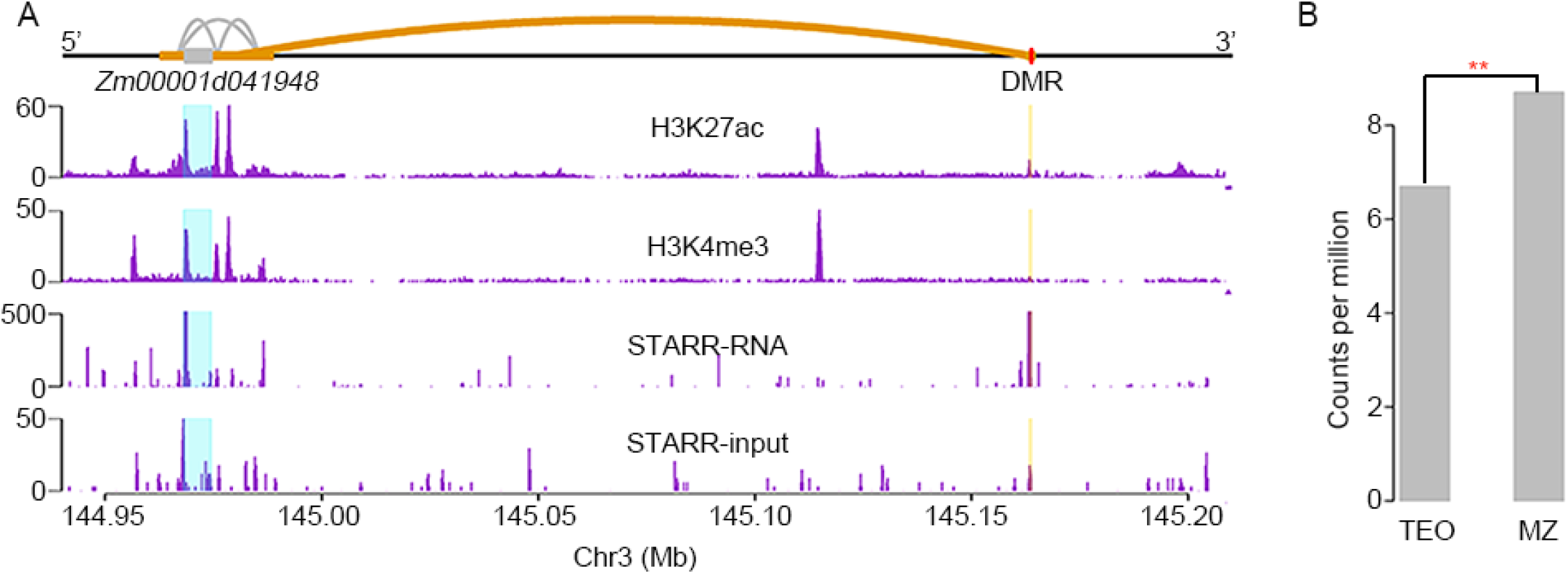
Interactive loops between a DMR and a gene model *Zm0001d041948.* (**A**) Chromatin interactions (the upper panel) and ChIP-Seq profiles (the lower panels) at gene *Zm00001d041948*. Gray and red boxes indicated the physical position of the gene model and the DMR. Gray and blue lines denoted the interactive loops. (**B**) Gene expression level of *Zm00001d041948* in teosinte and modern maize (Binomial test, P-value = 4.3e^-3^).

**Fig. S18.**
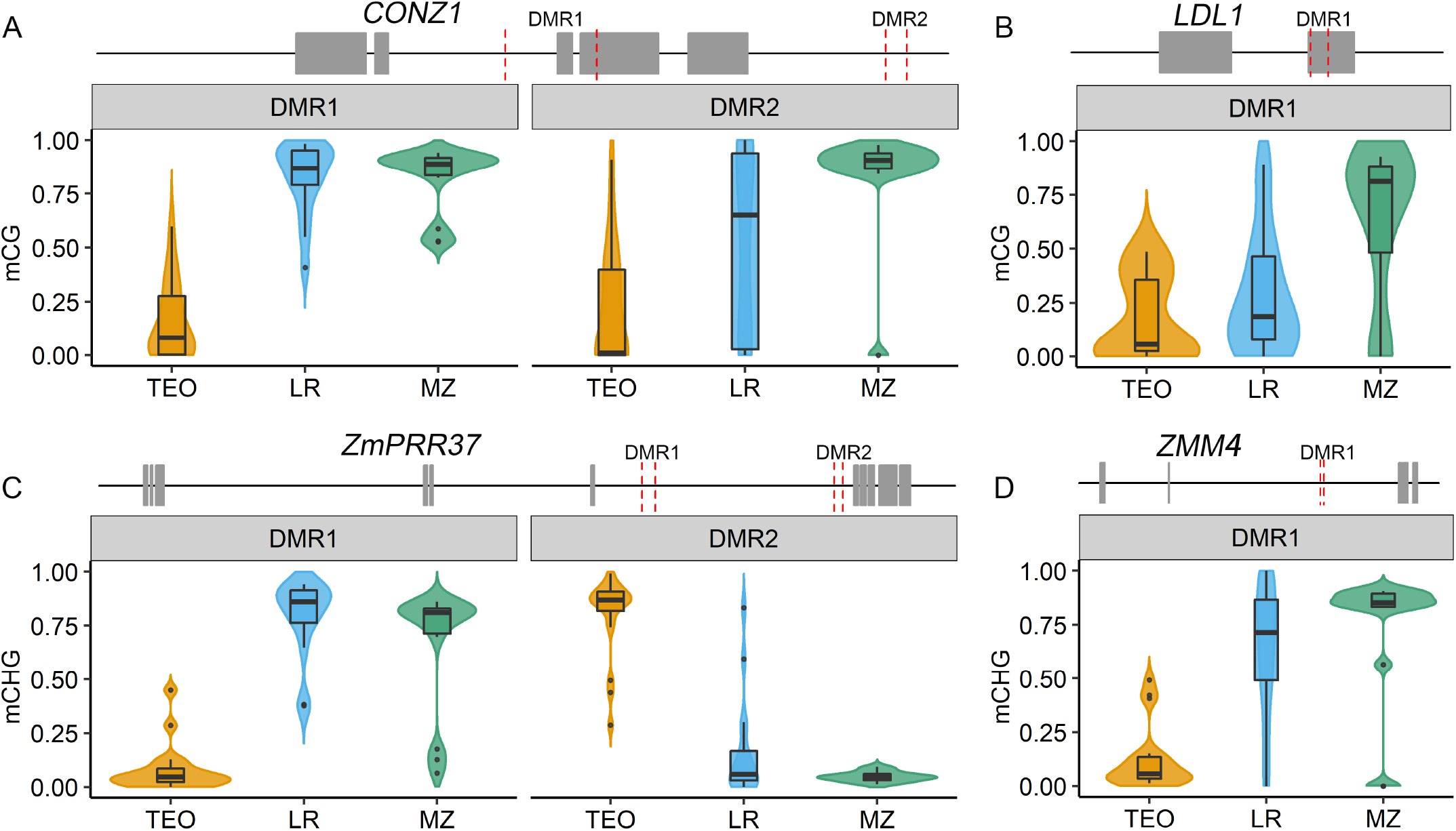
Teosinte-maize DMRs located at flowering time genes. Distribution of methylation level within DMRs that located at *CONZ1* (**A**), *LDL1* (**B**), *ZmPRR37* (**C**) and *ZMM4* (**D**) in teosinte (TEO), landrace (LR), and modern maize (MZ). Two nearby vertical dashed red lines on the gene model indicated a teosinte-maize DMR.

**Fig. S19.**
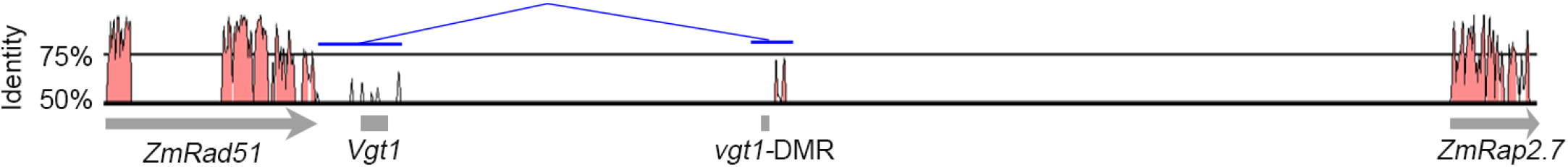
CNSs between the maize and sorghum orthologous around vgt1-DMR. Sequence identity of the maize sequence spanning *Vgt1* and the two proximal genes with corresponding sorghum sequences. Red peaks denote CNSs identified using a window size of 100 bp. The thin blue lines indicate physical interaction between two anchor sequences (thick blue lines).

**Fig. S20.**
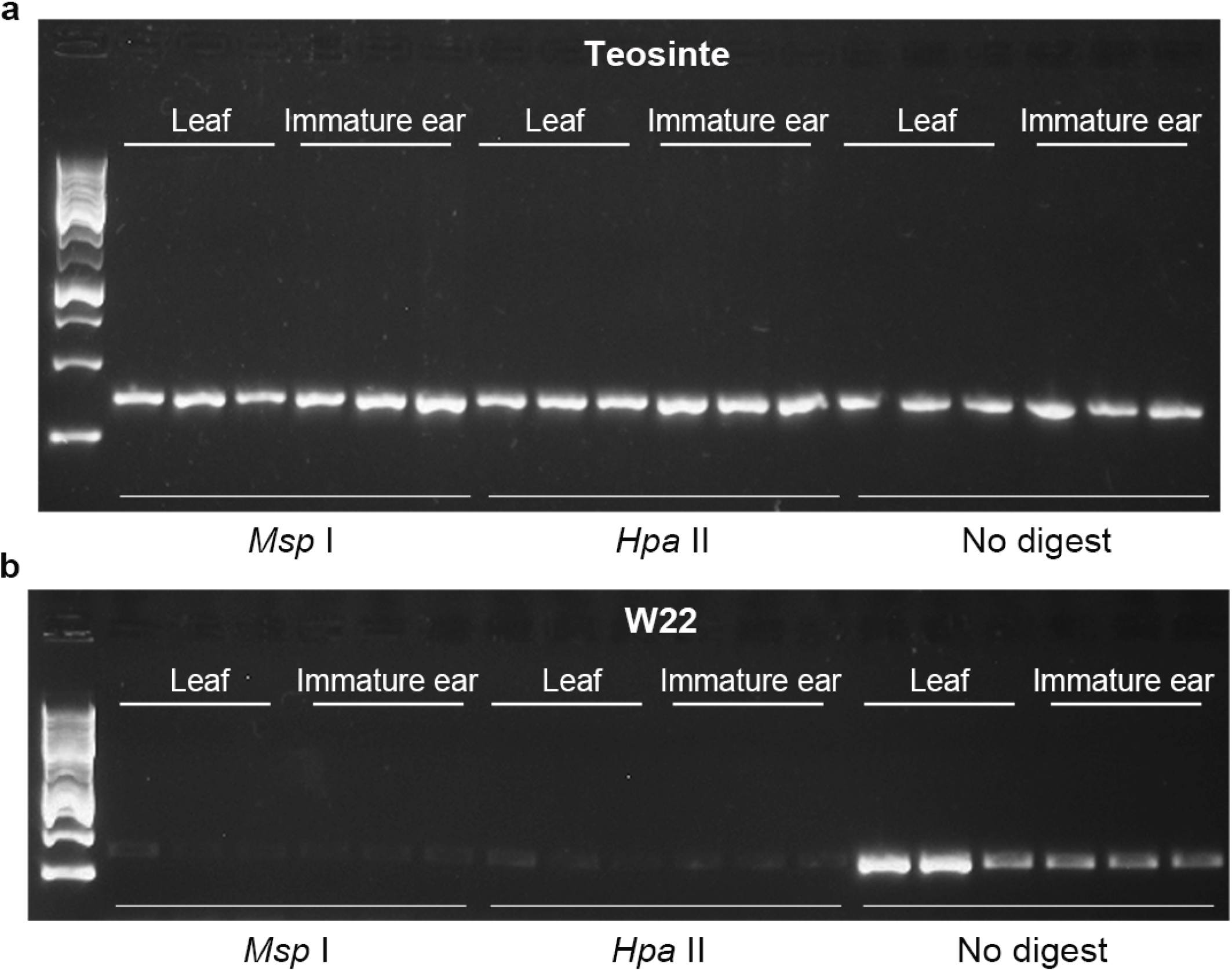
Experimental validation of the tb1-DMR using Chop-PCR. Chop–PCR analysis of *tb1*-DMR in different tissues of W22 (**A**) and teosinte 8759 (**B**). Failure to detect a PCR product reflects the loss of DNA methylation. CG methylation was detected using HpaII and CHG methylation was detected using MspI. Three independent biological replicates are shown, each with three technical replicates. No digested DNA is used as a control.

**Fig. S21.**
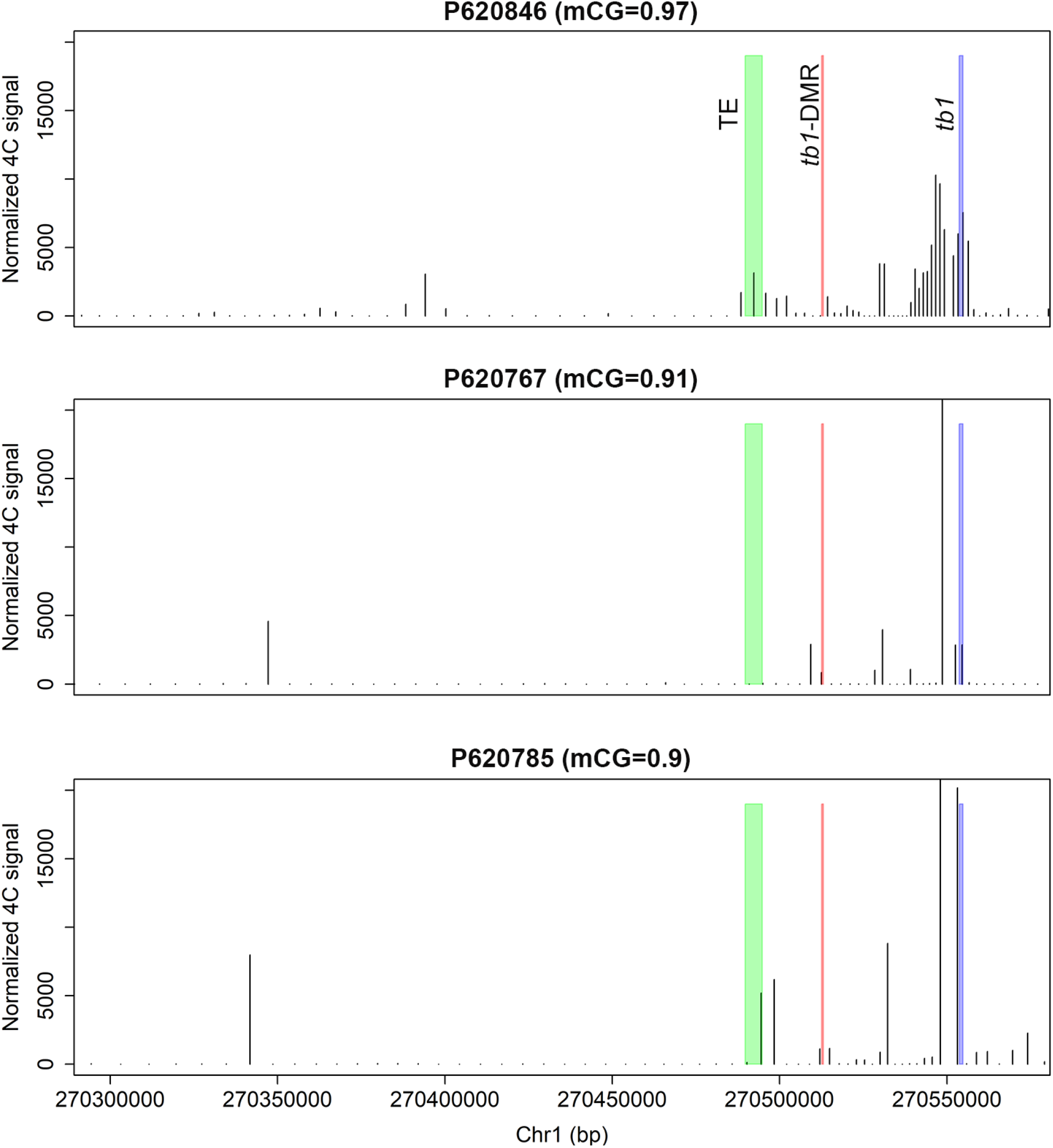
The 4C-seq results of regions interacted with *tb1* gene in tb1-DMR hypermethylated landrace samples. The titles show the CG methylation levels of the tb1-DMR. The green rectangle indicates hopscotch TE; the red rectangle indicates tb1-DMR; and the blue rectangle indicates *tb1* gene.

**Fig. S22.**
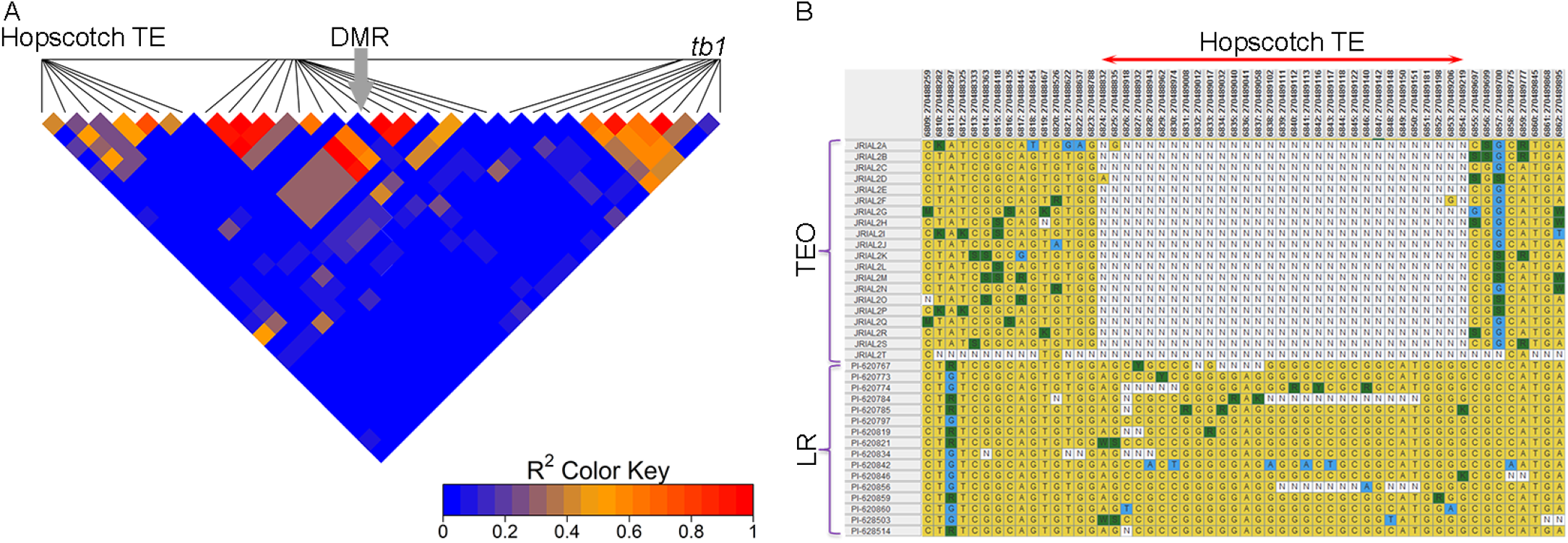
The linkage disequilibrium (LD) analysis among the Hopscotch transposon, tb1-DMR, and *tb1* gene. (**A**) The LD heatmap using 17 landraces segregating at the tb1-DMR locus. The arrow indicated the position of the tb1-DMR. LD analysis was performed using SNPs (coded with 0, 1, and 2) called from the WGS data and the mCG level of the *tb1-DMR.* The horizontal black line indicates the physical positions of the SNPs. (**B**) The SNP genotypes of teosinte and landrace around the Hopscotch transposon insertion region.

**Fig. S23.**
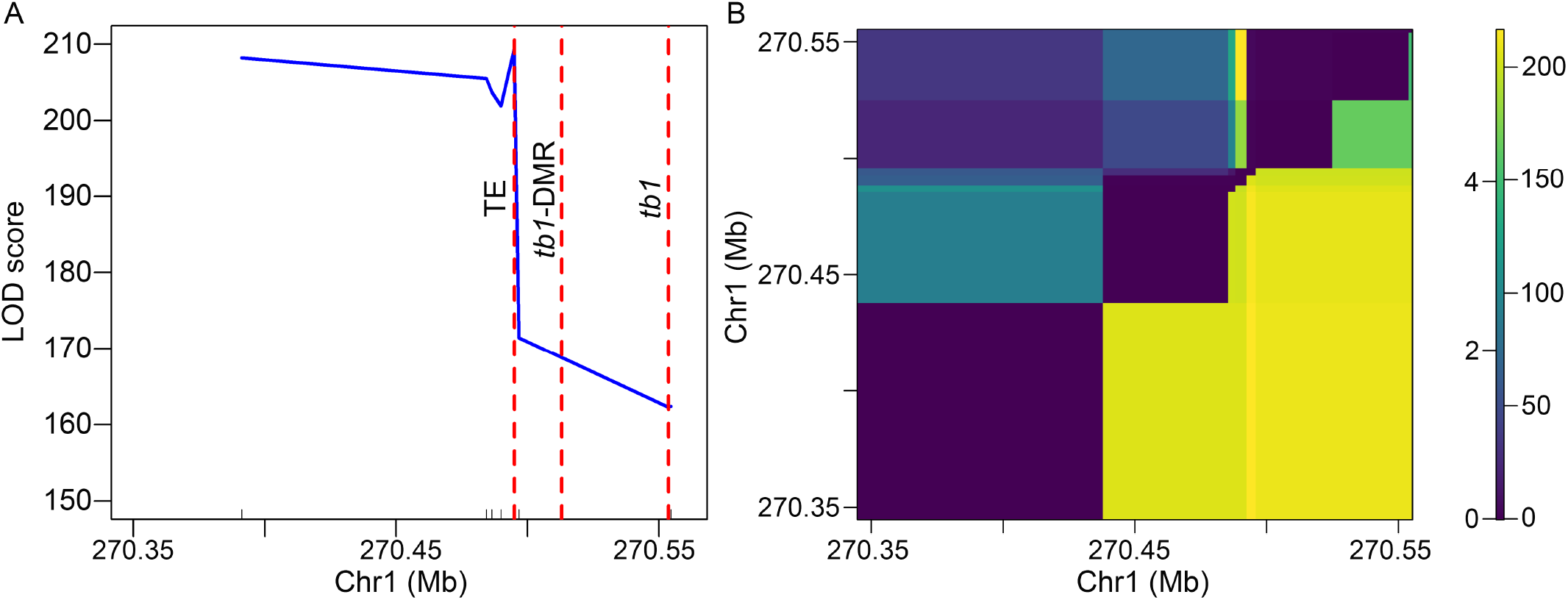
The QTL scannings for the tillering phenotype around tb1 locus. (**A**) The conventional single-QTL mapping result. Ticks above the x-axis indicated the physical positions of the markers. (**B**) Two-dimensional QTL scanning result. The lower triangle denoted the LOD scores for the joint two-locus; the upper left triangle denoted the LOD score for the epistasis of the two loci. The color scale on the right indicated LOD scores for the joint two-locus (right) and epistasis (left), separately.

**Fig. S24.**
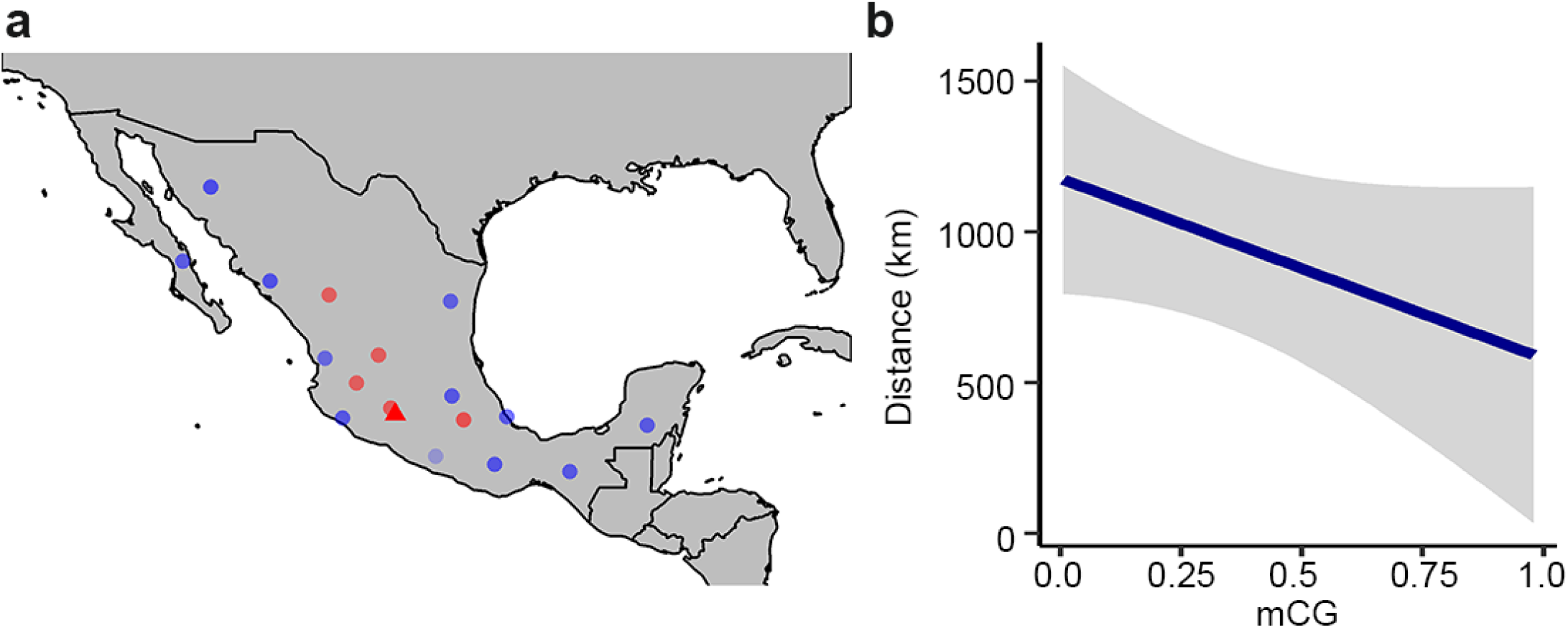
Correlation analysis between geographical distributions of landrace samples and their CG methylation level at tb1-DMR. (**A**) Geographical distributions of the teosinte (triangle) and landrace (points) samples. Red denotes highly methylated and blue denotes lowly methylated samples in the ŕb?-DMR. (**B**) Level of mCG correlated with distance to the origin (Balsas River Valley) of maize, with grey error bands marking the 95% confidence interval.

